# Gene–environment interactions govern early regeneration in fir and beech: evidence from participatory provenance trials across Europe

**DOI:** 10.64898/2026.01.29.702314

**Authors:** Katalin Csilléry, Justine Charlet de Sauvage, Madleina Caduff, Johannes Alt, Marjorie Bison, Mert Celik, Nicole Ponta, Daniel Wegmann

## Abstract

- Forest regeneration is shaped by strong demographic filters during germination and early establishment, yet the relative contributions of provenance and environment remain poorly resolved under natural conditions.
- We combined climate-chamber trials with a continental-scale, field-based citizen science seed-sowing experiment to quantify germination, early phenological development, and three-year post-germination survival fir (*Abies* spp.) and beech (*Fagus* spp.) provenances across Europe. Using mixed-effects models and a state-based Markov framework to reconstruct phenological trajectories, we quantified successive demographic filters from seed to established seedling.
- Germination was structured by genotype-by-environment interactions (*G × E*) in both genera, but with contrasting expression: fir showed broadly parallel environmental responses among provenances and partial concordance between climate-chamber and field, whereas beech showed strong provenance-by-environment responses and weak predictability from controlled conditions. Seed weight was a key axis in beech (lighter seeds germinated more and developed faster), but confounded with geographic differentiation in fir.
- In contrast to germination, post-establishment survival was dominated by site-level environmental filtering with limited consistent provenance effects, revealing a life-stage shift from fine-scale *G × E* during germination and early development to broader environmental control of survival—an important consideration for regeneration—based management and assisted migration.

## Introduction

Forest regeneration depends on adult tree reproduction - particularly seed production - as well as successful seed germination and seedling establishment [Donohue et al., 2010, Baskin and Baskin, 2000]. Despite the importance of these processes, the demographic, selective, and environmental factors that drive them remain poorly understood. Although heritable trait variation and adaptive differentiation between populations are common for growth traits in young adult trees [Savolainen et al., 2007, Alberto et al., 2013], the relative contributions of genetic, maternal and environmental effects to germination and early developmental traits are largely unknown [Manzanedo et al., 2018], especially under natural conditions [e.g., Muffler et al., 2021]. In the context of climate change, maladaptation and regeneration failure are becoming increasingly frequent in forest tree populations [Isaac-Renton et al., 2018], underscoring the urgency of assessing their adaptive potential.

The germination capacity of temperate forest tree species has been extensively studied in controlled laboratory conditions, often using treatments such as elevated temperature, reduced duration and various substrates during stratification to break dormancy, as well as different light regimes [e.g., Nawrot-Chorabik et al., 2021, Li et al., 1994, Escudero et al., 2002]. Wang et al. [2012], for instance, studied 86 subalpine forest tree species and found that while median germination time (*t*_50_) increased with seed mass across species, it was not consistently correlations with any broad environmental gradient. In a global meta-analysis of 63 tree species exposed to at least two temperature treatments in growth chambers, Vicente and Benito Garzón [2024] found that warming generally accelerated the timing of germination (reduced *t*_50_) more consistently across populations and biomes than it increased germination rates, and that responses were modulated by temperature at seed origin (e.g., warmer provenances exhibited larger reductions in germination time under elevated temperatures) [Vicente and Benito Garzón, 2024]. Finally, several authors have investigated the thermal optimum and thermal memory of tree seeds [e.g., Fernández-Pascual et al., 2019]. In conifers, a temperature optimum for germination is often observed, and interactions with moisture availability can further modulate both germination rate and success [Petrie et al., 2016, Hsu et al., 2024]. Finally, forest seed companies and national forest services also routinely conduct fast-germination tests [Bonner, 1998]. While these experiments are rarely published in academic journals and often follow protocols that do not always reflect natural regeneration conditions, they do provide insight into the intrinsic and environmental drivers of germination responses.

The germination capacity and early seedling development of forest tree species are also commonly assessed in seed nurseries and provenance trials. However, nursery-phase traits are typically recorded only until seedlings are transplanted to open-field sites [e.g., Frank et al., 2017], limiting their relevance for understanding the drivers of germination and establishment under natural conditions. Consequently, most available knowledge derives from observational studies of natural regeneration in the field [Redmond et al., 2018], particularly following natural disturbances [Dulamsuren et al., 2013, Moser et al., 2010, 2025, Valkonen and Maguire, 2005].

These studies consistently show that seedling emergence and early survival strongly depend on water availability and soil moisture and are therefore positively associated with canopy cover, cooler, wetter local climates, and the absence of summer drought. As a result, regeneration outside closed forests or after large disturbances is often restricted to exceptionally moist years or to pioneer species, often leading to long-lasting shifts in species composition [Valkonen and Maguire, 2005, Moser et al., 2010]. Despite this extensive observational evidence, experimental tests of germination and early establishment drivers under natural field conditions remain rare, mainly due to seed predation and drought-induced mortality [Walck et al., 2011]. To date, a continental-scale open-field transplant experiment explicitly targeting seed germination and early establishment has only been conducted for European beech [Muffler et al., 2021]. This study demonstrated that although germination success increased under warmer conditions, subsequent seedling establishment and survival declined in warmer and drier environments, highlighting a decoupling between germination and recruitment success [Muffler et al., 2021]. Conducting comparable experiments in additional species would be highly valuable to identify the environmental drivers of early regeneration stages and to complement the extensive experimental literature on bud break, chilling, forcing, and photoperiod responses in later life stages [e.g., Ma et al., 2021].

Beyond the limited number of experimental field studies, a further major challenge is that most research on tree regeneration has been conducted at the species level, whereas provenance-level variation in germination and early establishment remains poorly characterized. While regeneration strategies along the pioneer–shade-tolerant spectrum are well documented [e.g., Poorter, 2007], much less is known about whether regeneration capacity varies among provenances within species, or whether sufficient adaptive genetic variation exists for these traits to respond to climate change. Theory and empirical evidence suggest that early life stages are key targets of selection in long-lived organisms: because trees reproduce over many years, selection at the adult stage may be relatively weak, whereas intense mortality during germination and seedling establishment can generate population differentiation in early traits [Donohue et al., 2010, Kuparinen et al., 2010, Chang-Yang et al., 2021]. Indeed, decades of common-garden experiments in forest trees have revealed substantial adaptive genetic differentiation (*Q*_*ST*_ ) between provenances for early growth and phenological traits [Alberto et al., 2013]. Whether similarly strong differentiation exists at even earlier life stages remains largely understudied, primarily due to the substantial logistical challenges posed by the large sample sizes required, especially since mortality rates far exceed those of experiments preceded by a nursery phase. In a Mediterranean transplant experiment using one-year-old nursery-grown silver fir seedlings, provenance effects on survival and growth were modest in magnitude, with a large proportion of variation observed among families of the same provenance, highlighting substantial standing variation and environmental or stochastic heterogeneity at early life stages [Latreille and Pichot, 2017]. In European beech, Muffler et al. [2021] found significant provenance differences in germination capacity that were positively associated with seed weight, but no consistent differentiation in subsequent early-life traits. Finally, in *Pinus sylvestris*,common-garden experiments across Europe revealed provenance differentiation in seed weight, emergence timing, and early growth rate along climatic gradients [Ramírez-Valiente et al., 2021]. Overall, these studies suggest that although provenance-level variation exists in germination and early establishment, its magnitude and direction are highly context dependent.

Building on the limited empirical evidence for provenance-level variation in germination and early establishment under natural conditions, we investigated the joint roles of seed provenance, climate, and soil in shaping germination and early seedling survival in two major Eurasian forest tree species complexes: a conifer *Abies spp*. (firs) and a broadleaf *Fagus spp*. (beeches). We combined climate-chamber germination experiments, designed to estimate baseline germination capacity and timing across provenances under standardized conditions, with a continental-scale experimental approach in which seeds were sown directly in forests across Europe and monitored by voluntary foresters [Bison et al., 2025]. This integrative design allowed us to contrast provenance effects expressed under controlled conditions with those emerging under natural environmental variability, and to quantify the timing and strength of successive demographic filters from seed to established seedling.

Eurasian firs and beeches represent contrasting early-life histories and opposite ends of the seed size - seed production trade-off [Qiu et al., 2022]. Silver fir (*Abies alba* Mill.) is a dominant conifer of European mountain forests, distributed from the Pyrenees to the Carpathians [Mauri et al., 2016], and is increasingly regarded as a promising future species under climate change due to its relatively high drought tolerance, physiological plasticity, and capacity to persist under changing environmental conditions [Pistone et al., 2025, Vitasse et al., 2019]. We additionally included its closely related species *A. nordmanniana* (Steven) Spach, native to the Black Sea and Caucasus regions [Caudullo and Tinner, 2016]. Firs produce relatively few, light, winged seeds adapted for wind dispersal; seeds exhibit physiological dormancy requiring cold–wet stratification, do not form persistent seed banks, and recruitment may be delayed through the formation of seedling banks [Wolf, 2003]. In contrast, European beech (*Fagus sylvatica* L.), one of the most widespread broadleaf tree species in Europe, is considered particularly vulnerable to ongoing climate change, especially to increasing drought frequency and heat extremes, as evidenced by recent large-scale mortality and growth declines [Obladen et al., 2021, Frei et al., 2022]. We also included *F. caspica* Denk & Grimm, a closely related species endemic to the Hyrcanian forests of Iran [Denk et al., 2024]. Beeches produce a large number of heavy seeds with limited dispersal, experience intense post-dispersal predation, and lack seed banking [Wolf, 2008].

Our field experiment relied on a citizen-science framework, in which volunteer foresters established and monitored standardized experimental gardens (100 m^2^ each) across a broad climatic gradient. The experimental concept is described in Bison et al. [2025] for which this experiment served as a pilot. We adopted a hypothesis-driven framework, predicting that germination and very early development are comparatively deterministic and primarily influenced by seed quality and provenance-related conditions, whereas post-germination survival is more stochastic and largely governed by local environmental conditions at the garden sites. Because observations were conducted at irregular intervals by volunteers, we developed a state-based Markov model to reconstruct individual seed and seedling trajectories while accounting for observational uncertainty, completing the use of standard statistical approaches.

## Material and methods

### Seed origins

We used seeds from 12 silver fir (*Abies alba*) and nine European beech (*Fagus sylvatica*) provenances spanning their natural distribution ranges, and one provenance each of *A. nordmanniana* and *Fagus caspica* (Fig. 1, Table 1). Seeds were purchased from local seed providers or supplied by research institutes. All seeds were harvested in 2020 to ensure homogeneity, except for the Caspian beech *F. caspica* seeds from Iran, which were harvested in 2021. Given the large geographic distance and lack of masting synchrony between regions, differences between Caspian and European beech were attributed to species differences rather than harvest year.

**Table 1.** Fir (*Abies*) and beech (*Fagus*) provenances used in this study. ID is the abbreviated name of the provenance used through this study that incorporates the species abbreviation (AA for *Abies alba*; AN for *A. nordmanniana*; FS for *Fagus sylvatica*; FC for *F. caspica*), the country, and a provenance number. Provenance is the name used by the seed provider or the name of the region/mountain range in the absence of it. Seed weight (SW, g per 100 seeds) and seed moisture content (SMC, %) are reported where available. Longitude (Lon), latitude (Lat), and elevation correspond to seed origin locations. Use indicates whether a provenance was included in climate chamber trials (CC), micro-garden field trials (MG), or both (CC/MG).

| ID | Country | Provenance | Lat (°N) | Lon (°E) | Elev. (m) | Source | SW | SMC | Use |
| --- | --- | --- | --- | --- | --- | --- | --- | --- | --- |
| AA_AT1 | Austria | Zwischenalp South | 47.292 | 15.190 | 900 | Austrian Research Centre BFW | NA | NA | MG |
| AA_DE1 | Germany | Traunstein | 47.797 | 12.810 | 875 | Bayerische Staatsforsten AöR | NA | NA | MG |
| AA_FR1 | France | Corse | 42.250 | 8.800 | 1100 | French National Forests Office | 4.66 | 6.43 | CC/MG |
| AA_FR2 | France | Haute Provence | 44.050 | 6.510 | 990 | French National Forests Office | 3.65 | 8.65 | CC/MG |
| AA_FR3 | France | Pyrenees | 42.853 | 1.676 | 1100 | French National Forests Office | 4.23 | 7.97 | CC/MG |
| AA_FR4 | France | Massif Central West | 45.900 | 3.450 | 475 | French National Forests Office | NA | NA | MG |
| AA_IT1 | Italy | Lavarone Trento | 45.957 | 11.299 | 1230 | NA | 3.87 | 9.18 | CC/MG |
| AA_PL1 | Poland | Sudety | 50.934 | 15.832 | 900 | The Kostrzyca Forest Gene Bank | 6.50 | 12.25 | CC/MG |
| AA_RO1 | Romania | SE Carpathians | 45.699 | 26.462 | 650 | Transilvania University of Braşov | 5.71 | 12.99 | CC/MG |
| AA_SK1 | Slovakia | North Region | 49.300 | 19.300 | 890 | LESY Slovenskej republiky | 4.41 | 9.64 | CC/MG |
| AA_SI1 | Slovenia | Pohorsko | 46.533 | 15.250 | 900 | Slovenia Forest Service | NA | NA | MG |
| AA_HR1 | Croatia | Crni Lazi | 45.570 | 14.613 | 914 | Croatian Forest Research Institute | 4.43 | 8.81 | CC/MG |
| AN_GE1 | Georgia | Ambrolauri | 42.514 | 43.146 | 1140 | HD Seed | 6.72 | 5.67 | CC/MG |

Table 1: Table 1 continued.
| ID | Country | Provenance | Lat (°N) | Lon (°E) | Elev. (m) | Source | SW | SMC | Use |
| --- | --- | --- | --- | --- | --- | --- | --- | --- | --- |
| FS_AT1 | Austria | Randgebirge South | 46.445 | 14.268 | 900 | Austrian Research Centre BFW | NA | NA | MG |
| FS_FR1 | France | Massif Armoricaïn | 48.370 | 0.100 | 260 | French National Forests Office | 24.86 | 14.70 | CC/MG |
| FS_FR2 | France | Massif Central North | 44.780 | 1.880 | 500 | French National Forests Office | NA | NA | MG |
| FS_IT1 | Italy | Passo Fittanze | 45.684 | 10.989 | 1370 | CNC Biodiversità Peri | 21.62 | 8.07 | CC/MG |
| FS_RO1 | Romania | SE Carpathians | 47.818 | 25.608 | 792 | NA | 26.72 | 14.71 | CC/MG |
| FS_SK1 | Slovakia | Middle Region | 49.000 | 19.950 | 845 | LESY Slovenskej republiky | 24.37 | 9.99 | CC/MG |
| FS_SI1 | Slovenia | Dinarsko | 45.600 | 15.050 | 782 | Slovenia Forest Service | 22.00 | 9.20 | CC/MG |
| FS_CH1 | Switzerland | Salenstein | 47.669 | 9.059 | 400 | Forstbaumschule Kressibucher AG | 26.98 | 14.68 | CC/MG |
| FC_IR1 | Iran | Alborz Mts. High | 36.304 | 52.143 | 1873 | Caspian Tree Seed Center | NA | NA | MG |
| FC_IR2 | Iran | Alborz Mts. Middle | 36.315 | 52.129 | 1442 | Caspian Tree Seed Center | 24.34 | 13.20 | CC/MG |

**Figure 1.**
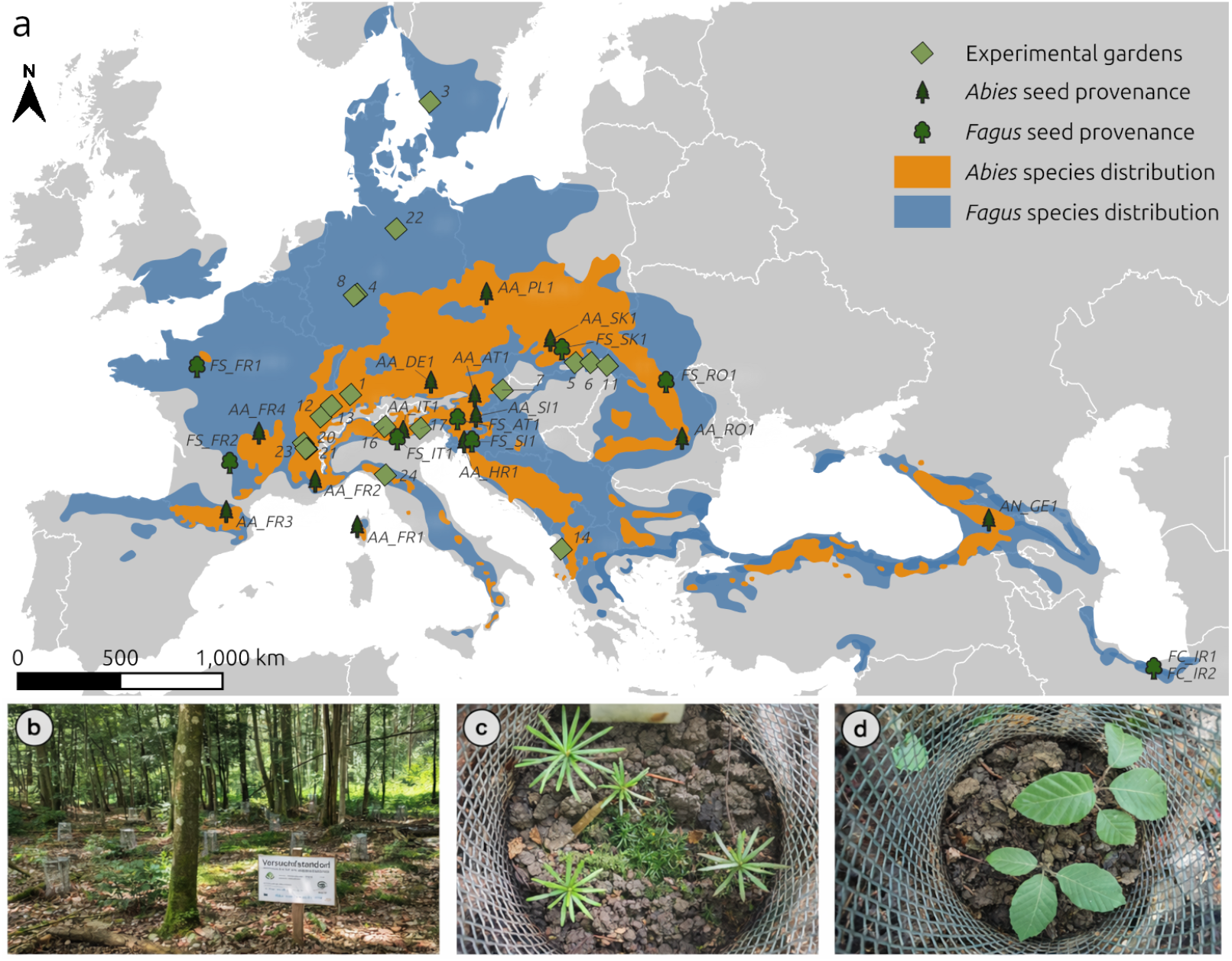
Study design with the location of fir (*Abies*) and beech (*Fagus*) seed provenances and the transplantation sites called micro-gardens. (a) Geographic distribution of seed provenances (Table 1) and micro-gardens (Table 2). Shaded areas indicate the natural distribution ranges of *Abies* (orange) and *Fagus* (blue) species complexes. (b) Example of a micro-garden established under forest canopy. (c) Seed protector containing emerging *Abies* seedlings. (d) Seed protector containing emerging *Fagus* seedlings. Photographs in panels (b–d): Justine Charlet de Sauvage.

Upon reception, seeds were stored at 5 *±* 0.5 ^*°*^C until the start of the experiments. For each provenance tested in the climate chamber trials, seed moisture content (SMC) was determined from 100 seeds following the low-temperature drying method recommended by the International Seed Testing Association (ISTA; Grabe, 1989) and described by Bezdeckova et al. [2014]. Fresh weight was measured using a precision balance (Mettler Toledo XS603S, 1 mg readability), after which seeds were dried for 17 *±* 1 h at 103 *±* 2 ^*°*^C. Seed moisture content was calculated as:

**Table 2.** Description of the micro-gardens. Coordinates and elevation represent average values across blocks within each garden. Garden 13 and 21 are reported as two separate entries corresponding to groups of blocks at contrasting elevations; therefore, they are treated as separate gardens. All other gardens contain four blocks in close proximity (A1, A2, F1, and F2). The sowing date is given in the format dd.mm.yyyy.

| Garden_ID | Country | Lat (°N) | Lon (°E) | Elevation (m) | Sowing date |
| --- | --- | --- | --- | --- | --- |
| 1 | Switzerland | 47.363 | 8.455 | 598 | 13.01.2022 |
| 3 | Sweden | 57.056 | 12.766 | 159 | 05.03.2022 |
| 4 | Germany | 50.931 | 8.783 | 348 | 04.03.2022 |
| 5 | Hungary | 48.521 | 20.672 | 223 | 18.02.2022 |
| 6 | Hungary | 48.523 | 21.524 | 368 | 24.02.2022 |
| 7 | Hungary | 47.529 | 16.707 | 289 | 22.02.2022 |
| 8 | Germany | 50.877 | 8.650 | 382 | 24.02.2022 |
| 11 | Ukraine | 48.421 | 22.421 | 107 | 22.02.2022 |
| 12 | Switzerland | 46.534 | 6.842 | 750 | 16.02.2022 |
| 13_F1 and 13_F2 | Switzerland | 46.956 | 7.415 | 601 | 01.04.2022 |
| 13_A1 and 13_A2 | Switzerland | 46.927 | 7.399 | 715 | 01.04.2022 |
| 14 | Albania | 41.360 | 19.910 | 1000 | 23.02.2022 |
| 16 | Italy | 46.172 | 10.326 | 748 | 22.02.2022 |
| 17 | Italy | 46.101 | 12.252 | 731 | 20.02.2022 |
| 20 | France | 45.537 | 5.886 | 624 | 11.04.2022 |
| 21_A1 and 21_F2 | France | 45.376 | 6.088 | 718 | 10.02.2022 |
| 21_A2 and 21_F1 | France | 45.280 | 6.064 | 1400 | 19.02.2022 |
| 22 | Germany | 53.114 | 10.925 | 103 | 05.02.2022 |
| 23 | France | 45.333 | 6.026 | 1056 | 18.02.2022 |
| 24 | Italy | 44.315 | 10.342 | 984 | 08.02.2022 |

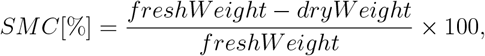

assuming that dried seeds had 0% moisture content.

In addition, 220 seeds per provenance were weighed before and after pre-treatment to estimate changes in fresh weight. Of these, 200 seeds were used in germination experiments, while 20 served as a buffer to account for potential losses during pre-treatment (e.g., fungal infestation or handling). Fresh weight measurements from the 100 seeds used for moisture determination and the 220 seeds assigned to pre-treatment were combined to estimate the weight of 1,000 seeds (SW, g; Table 1, used here as a proxy for seed mass).

### Climate chamber trials

Baseline germination and early seedling development were assessed in climate chamber experiments. Due to space limitations, experiments were conducted in two consecutive cycles, each including up to eight provenances (the first cycle was published in Alt 2023). In total, nine *Abies* and seven *Fagus* provenances were tested (Table 1), selected based on seed availability and geographic representation. *Abies* seeds were cold-stratified for four weeks at 5 *±* 0.5 ^*°*^C on moist blotting paper, whereas *Fagus* seeds were soaked in rainwater for 48 h at the same temperature. For each provenance, 200 seeds were tested, divided equally between two watering treatments (wet vs. dry). In each treatment, 100 seeds were sown in a 51 *×* 31 cm plastic tray filled with a 1:1 mixture of sand and peat (Fig. S1).

Trays were watered once (dry treatment) or twice (wet treatment) per week until substrate saturation and were repositioned twice weekly within the climate chamber to minimize spatial heterogeneity. A photoperiod of 8 h light (20 *×* 1280 W lamps) and 16 h darkness was applied.For the first 58 days, temperatures were set to 15 *±* 0.5 ^*°*^C during the day and 5 *±* 0.5 ^*°*^Cat night, followed by a warming phase of 14 days at 20/10 *±* 0.5 ^*°*^C (day/night) to promote germination of remaining dormant seeds. Relative humidity was maintained at 70 *±* 5%.

Seed germination was assessed twice weekly by visual inspection, and trays were photographed at each observation. Seedlings were classified into four phenological stages for fir and five stages for beech species (Fig. S2). At the end of the experiment, all remaining seeds were recovered from the substrate and examined for radicle emergence. Because numerous viable but ungerminated seeds were detected at the end of the first cycle, the second cycle was extended by 32 days.

### Field trials

In summer and autumn 2021, 18 volunteer foresters were recruited across Europe to establish field experiments, hereafter referred to as micro-gardens, directly under the forest canopy on sites they owned and/or managed (Table 2, Fig. 1a, Methods S1). Each micro-garden covered approximately 100 m^2^, contained 100 seeding spots, and was divided into four blocks, two per genus (A1 and A2 for *Abies*; F1 and F2 for *Fagus*; see Supplementary Document S1 for protocols) with 25 seeding spots per block. At each seeding spot, 10 seeds from a single provenance were placed on the ground and covered with a thin layer of local soil and litter. A purpose-designed seed protector was installed at each spot to reduce seed predation and prevent the dispersal of seeds from other provenances or genera (Fig. 1b-d). Each provenance was replicated five times per micro-garden. Seeds were sown in winter 2021–2022 to allow natural stratification under local climatic conditions (Table 2).

Monitoring was conducted according to volunteer availability, with a target frequency of weekly visits during the main germination period (at least six visits between April and May) and monthly visits from June to October (see Supplementary Document S1). Germination phenology was recorded in spring 2022 using the same criteria as in the climate-chamber experiment. Bud-break phenology was monitored in spring 2023 and 2024, and seedling survival was assessed during summer and autumn in each of the three study years. Seeds and seedlings were monitored at the seed-protector level, which remained in place throughout the study period (Fig. 1b–d). Observations were recorded using a custom web-based form implemented in ODK Collect [Ouma et al., 2019] (Dataset S1).

### Climate and soil data

We used climate and soil data to (i) characterize the long-term environments at seed source locations (provenance environments), (ii) describe the environmental conditions experienced by seeds and seedlings at the micro-garden sites (garden environments), and (iii) derive thermal indices used to parameterize stage transitions in the Markov model. For the first two groups of variables, see Table S1 for details.

First, to characterize the long-term climate at seed source locations, we extracted monthly temperature and precipitation time series from ClimateDT (1 km resolution) [Marchi et al., 2024]. We summarized provenance climate using monthly mean temperature and monthly precipitation totals across the 12 months of the year, and annual temperature and annual precipitation across the period 1901–1980; a baseline period chosen to reduce the influence of recent anthropogenic warming on provenance characterization (Supplementary Data S2). Second, daily temperature and precipitation at micro-garden sites were extracted from the CHELSA v2.1 daily product (30 arcsec, *∼*1 km), which provides global daily surface variables including near-surface air temperature and precipitation [Karger et al., 2021, 2017]. We extracted daily data from September 2021 (to cover the pre-sowing period) through October 2022 (covering the first growing season), at the block level within each micro-garden. CHELSA-daily is released as an actively updated dataset and may contain gaps for the most recent period [Karger et al., 2021]. If a single day of daily mean temperature was missing, we gap-filled it using linear interpolation based on the means of the preceding and following days. Blocks of a micro-garden typically fell within the same CHELSA grid cell. However, blocks of micro-gardens 13 and 21 were established in spatially separated locations with contrasting environments. Therefore, climate data were extracted at the block level, and these locations were treated as separate garden units in the mixed-effects analyses. For garden 13, this subdivision is not apparent in the results because the two contrasting environments correspond to the two genus blocks (Table 2).

Third, to model seed-to-seedling and seedling phenological stage transitions, we derived indices from the daily CHELSA data. Growing degree-days (GDD) were calculated as the cumulative sum of daily mean temperature above a base threshold of 5 ^*°*^C, starting on 1 January of each year. We used this index to parameterize development (“growth”) as a function of accumulated thermal time in the Markov model (see below). For other analysis, we computed a cumulative GDD up to 1 July 2022 (i.e. removing the time aspect) and summarized spring conditions using mean temperature and precipitation sums for selected months during the germination and early establishment period (November 2021–June 2022). Additionally, chilling days were computed as the number of days since sowing with daily mean temperature below 5 ^*°*^C. For garden 20 (sown on 11 April 2022 due to logistical difficulties), seeds did not experience chilling in the field. To align pre-sowing cold exposure with the other gardens, *Abies* seeds were cold-stratified at 5 ^*°*^C on moist paper from 19 February 2022 (51 days) and *Fagus* seeds were soaked at 5 ^*°*^C for 48 hours before sowing.

Finally, soil properties at provenance and micro-garden locations were extracted from SoilGrids (global 250 m soil property maps) [de Sousa et al., 2021]. We used a single depth interval (15–30 cm) to obtain a balanced set of climate-soil predictors while targeting the upper mineral soil layer relevant for early root development and soil water availability relevant for forest regeneration and seedling establishment [Puhlick et al., 2021, Socha et al., 2022, Csilléry et al., 2020b]. The candidate soil predictors were: bulk density, cation exchange capacity, coarse fragments, clay, sand, and silt fractions, soil organic carbon, total nitrogen, organic carbon density, and soil pH in water, extracted for both provenances and gardens.

#### 0.1 Germination metrics and survival analysis from the climate chamber trials

To explore the climate chamber data, we first used the R package `germinationmetrics` to calculate the final germination percentage (function `GermPercent`), the median germination time (function t50) using the formula of Coolbear et al. [1984], and measures of germination uniformity (function `CVGermTime` ) and synchrony (function `GermUncertainty and GermSynchrony` ), and correlated these with geography and seed traits. To compare the provenances, we estimated the survival probability *S*(*t*) over time using the Kaplan-Meier (KM) method implemented in the `survfit` function of the R package survival [Therneau, 2024]. We then used the Cox proportional hazards model implemented in the `coxph` function to test differences among provenances and the effect of treatment on germination probability, and, in a separate model, to evaluate the effects of longitude, seed weight, and moisture content. We ran separate models for the two sets of covariates because the number of provenances did not allow us to test all simultaneously. We also ran separate models for the two genera. Effects were plotted using the `forest_model` function of the R package `forestmodel` . To allow comparisons across provenances from different cycles, we considered only the first 72 days of each cycle.

### Hidden Markov model for germination and development

We developed a Markov model to estimate the germination rate (*g*), the development speed (*δ*), and the probability to survive until a given time (*P* (*x*)) for each unique combination of provenance and micro-garden environment. Observations of the number of alive seedlings and their phenological stage formed a time series for each seed protector, our observation unit (Fig. 1c and d). The observed counts of alive seedlings at any particular seed protector arose from a combination of three categories of individuals: (i) viable seeds that are dormant and have not yet germinated, (ii) seeds that have germinated into currently growing seedlings, and (iii) non-viable seeds or a previously germinated seedling that have died. The probability of germination depended on the growing degree days (GDD) at each micro-garden location, modeled as a logistic function. Then, the seedlings were assumed to grow linearly after germination, with growth rates specific to their phenological stage (Fig. S2). The probability of death for a currently growing seedling was determined by its current size and modeled to be highest at germination, then to decline exponentially. The relationship between growth and phenological stages was modeled using Gamma distributions and learned from climate chamber data. Since the protocol did not track individual seeds but only counts per seed protector, we used importance sampling to integrate over all possible seed trajectories that could give rise to the observed counts. Briefly, we simulated germination dates and death intervals by randomly selecting germination dates that precede death intervals, subject to the model’s constraints (see Methods S2 for a detailed description of the full model).

We applied the Markov model described above to both the climate chamber and micro-garden data to estimate *g, δ*, and *P* (*x*). In the climate chamber, parameter estimates were obtained for each provenance on the pooled data of the dry and wet treatments. In the field trials, parameter estimates were obtained for each combination of provenance and micro-garden location; blocks were pooled (except for IDs 13 and 21; Table 2) to increase the available data for parameter estimation, and we accounted for each garden’s specific GDD. We restricted the analysis to the first year (2022) data, which had the largest number of observations and included the germination phase, enabling comparison with the parameter estimates from the climate chambers. Since the much denser observation intervals in the climate chamber allowed us to observe the transitions between phenological stages and the time spent in each stage (Fig. S2), we used the emission probabilities estimated from the climate chamber data also for the field trials.

### Mixed effects models for the field trials

While we used the Markov model to explore the role of gene-environment interactions in germination rate and phenology, we could obtain only one parameter value per provenance-environment combination, limiting our ability to identify abiotic selection filters during regeneration. We therefore complemented these estimates with an approximate estimate of the germination rate per seed protector for use in a mixed-effects model (see below). Remember that field observations were conducted at irregular intervals (see Fig. S10) and on groups of 10 individuals each. It took approximately four weeks from the first germination to the last germination across all gardens and provenances in both fir and beech (Fig. S11 and S12), respectively. Yet, the average length of observation intervals was on average 12.5 days (SD: 8.2 days) across all gardens during the first spring of the experiment (between 1 April and 30 June 2022). As a consequence, changes in the number of observed seedlings between two observation dates could reflect newly germinated individuals, mortality, or both. To disentangle these processes, we used information on seedling phenological stages to infer the most likely transitions between consecutive censuses. Between two consecutive observations (*t* and *t* + 1), seedlings observed at *t* + 1 were matched to those observed at *t* based on their phenological stage progression, assuming that individuals in more advanced stages at time *t* had a higher probability of persisting to *t* + 1 than those in earlier stages. When multiple assignments were possible, matches were therefore resolved by preferentially attributing surviving individuals at *t* + 1 to the most advanced compatible stage at *t*. After accounting for these stage-based matches, the number of newly germinated seedlings at time *t* + 1 was defined as the number of observed individuals that could not be assigned to any seedling present at time *t*. This procedure was applied to all observations up to 31 October 2022. Then, we calculated the cumulative maximum germination rate across all observation dates up to 31 June 2022, thereby eliminating the time dimension from our high-dimensional, sparse matrix of observations and obtaining a response variable that could be modeled with a negative binomial distribution (`family = asr negative.binomial` ).

We fitted linear mixed-effects models using ASReml-R version 4.2 [Butler et al., 2023] to quantify the influence of geographic, environmental, and seed traits on germination. A separate modeling procedure was applied for fir and beech. First, to reduce the large set of environmental covariates (71 variables) and avoid overfitting, we applied a LASSO regression [Tibshirani, 1996] using the `glmnet` R package [Friedman et al., 2010]. All predictors were standardized prior to the analysis, and the penalization parameter *λ* was chosen by ten-fold cross-validation (function `cv.glmnet` ). We retained 19 (fir) and 13 (beech) variables with non-zero coefficients at *λ*_min_ as associated with germination (Fig. S4). We then performed clustering of the variables based on their correlations and selected variables relevant to both genera and those that were integrative, such as the mean across several months, rather than monthly variables or those that were easier to interpret. The final set of fixed effects included the seed traits (seed weight and moisture content) and the selected environmental variables. All continuous predictors were standardized before analysis. We included the following random effects: provenance identity, garden identity, and the provenance-by-garden interaction. All variance components were expressed as proportions of the total variance. All models converged and produced a satisfactory fit to the data as evaluated by diagnostic plots (Fig. S14).

Finally, seedling survival across the three growing seasons was analyzed at the cohort level (unique garden-provenance combinations) using a binomial generalized linear mixed model with logit link. The total number of seedlings for each cohort was taken as the number observed at the peak germination date, i.e., when the number of seedlings was highest. Initial cohort size (*N*_0_) was defined as the maximum aggregated seedling count observed during the establishment year (2022). Cohorts that did not germinate were excluded from this analysis. To separate the post-establishment survival from germination dynamics, we defined the start of the survival period as the first census date at which a cohort reached its maximum abundance. All observations prior to this date were excluded. Survival time was then expressed as the number of days since the cohort peak for each observation. Although the calendar date of the cohort peak varied among cohorts (standard deviations of 22 and 28 days for fir and beech, respectively), this variation did not explain additional survival differences once survival time was expressed relative to the cohort peak (not shown). Survival time was grouped into ecologically meaningful temporal bins that represent major seasonal abiotic filters acting on seedlings in temperate environments. They approximately corresponded to: early summer establishment, late summer drought, autumn transition, winter freezing period, spring thaw, and subsequent annual cycles. Bin boundaries were chosen to balance biological interpretability with adequate sample size per bin (at least 20 observations). Seedling survival was analyzed using a binomial generalized linear mixed model using a logit link fitted in ASReml. For each cohort and census date, the number of surviving seedlings was modeled as a binomial response with *N*_0_ as the denominator. The fixed-effect structure included the categorical temporal bin factor described above, capturing seasonal variation in survival probability. To account for hierarchical structure and repeated measurements, garden identity and provenance (ID) were included as random intercepts. Further, to accommodate extra-binomial variation arising from repeated counts, observer effects, or micro-site heterogeneity, an observation-level random effect (OLRE) was included. The predicted survival probabilities were obtained for each temporal bin at the genus level (separate models were run for fir and beech) and for field gardens. Garden-specific predictions were used to visualize heterogeneity between micro-gardens, while population-level predictions and 95% confidence intervals were used to summarize overall survival patterns of each genus.

## Results

### Germination and development in the climate chambers

Two beech provenances did not germinate at all (FS CH1 and FS RO1) in the climate chambers, and FS IT1 showed very low germination (2% and 3% in the dry and moist treatments, respectively; Fig. S5, Table 1). These provenances were excluded from further analyses because their failure to germinate could not be attributed unambiguously to biological rather than storage-related causes. In addition, the number of germinated seeds for FS IT1 was too small to reliably calculate germination metrics.

The length of the observation cycle had a predictable and consistent effect on germination parameters across provenances (Fig. S8). Root inspections at the end of the experiment indicated slightly higher germination counts than visual assessments alone (Fig. S5). All provenances tested in the second, 32-day-longer cycle reached a germination plateau shortly after 72 days, indicating that a 72-day observation window was sufficient to estimate germination parameters for both genera. Consequently, observations were truncated at 72 days for subsequent analyses, except for Kaplan–Meier curves, which are shown for the full duration.

Across both genera, germination metrics spanned similar ranges but exhibited pronounced within-genus and among-provenance variation (Fig. 2a,d; Fig. S5; Table S1). In both genera, eastern provenances tended to germinate more successfully, and provenances with lower germination rates also tended to germinate more slowly and less synchronously (Fig. S7).

**Figure 2.**
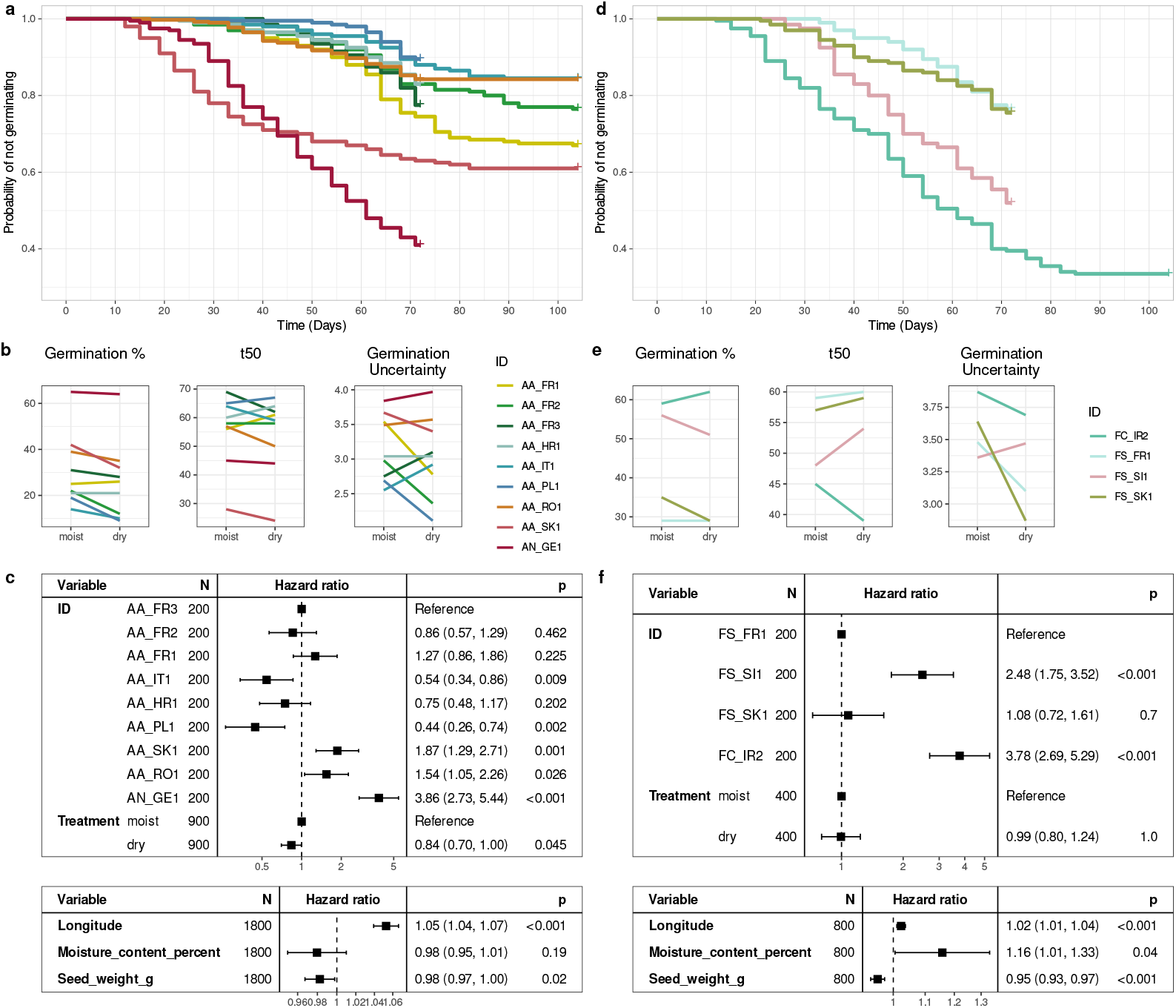
Germination of fir and beech seedlings in controlled climate chamber experiments. (a, d) Kaplan-Meier survival curves of fir and beech provenances, respectively. (b, e) The effect of moisture treatment on germination metrics for fir and beech, respectively. (c, f) Hazard ratios estimated from two Cox proportional hazards regression models for provenance differences and treatment effects (top panel), and for longitude, latitude, and seed traits (bottom panel) for fir and beech provenances, respectively.

Markov model parameter estimates derived from climate chamber observations closely matched simple germination metrics, validating the modelling framework used for the field data. In fir and beech, the estimated germination rate (*g*) was strongly correlated with the proportion of germinated seeds (r = 0.95 and 0.98, respectively; Fig. S7). In contrast, the development speed parameter (*δ*) showed only weak associations with germination metrics, indicating partial decoupling between germination and early phenological development. Within this general pattern, substantial provenance-level differences were detected. In fir, Nordmann fir showed the highest germination percentage (64%), exceeding those of the next-best provenances from Slovakia (AA SK1, 40%) and Romania (AA RO1, 37%) by more than 30 percentage points. In beech, the highest germination was observed for *Fagus caspica* from the Alborz Mountains (FC IR2, 59%), closely followed by the Dinaric provenance from Slovenia (FS SI1, 54%).

Cox proportional hazards models also confirmed strong provenance effects on germination timing in both genera (Fig. 2c,f). In fir, increased moisture significantly reduced germination probability, whereas no such effect was detected in beech. Germination metrics further revealed provenance-specific responses to water availability (Fig. 2b,e): fir provenances generally germinated less under dry conditions, while Iranian beech showed higher germination under dry treatment. Germination speed, uniformity, and synchrony responded less predictably, indicating complex provenance-by-environment interactions. Models incorporating geographic origin and seed traits revealed a weak but significant east–west gradient in germination probability in both genera. In beech, higher seed moisture content increased germination probability, while in both genera lighter seeds germinated more successfully (Fig. S7b). Finally, seed weight influenced early post-germination development under controlled conditions. In both species, development speed (*δ*) was negatively correlated with seed weight (fir: r = –0.71, *P* = 0.031; beech: r = –0.97, *P* = 0.027), indicating that lighter seeds progressed more rapidly through early developmental stages.

### Germination and development in the field trials

Field-based estimates of germination rates (*g*) demonstrated that the micro-garden environment and provenance co-determined seedling establishment in both genera (Fig. 3c,d). In fir, some provenances, most notably from the southeastern Romanian Carpathians (AA RO1) and Nordmann fir from Georgia (AN GE1), showed consistently higher germination across gardens; however, their mean germination rates remained comparable to, or lower than, the maximum germination achieved by other provenances in individual gardens. Similarly in beech, the Slovenian Dinaric provenance (FS SI1) exhibited exceptionally high germination, exceeding 50% in two gardens and showing a mean field germination rate (21.8%) more than twice that of the next best provenance (FS IT1, *g* = 9.4%), but other provenances achieved higher germination rates in several individual gardens.

**Figure 3.**
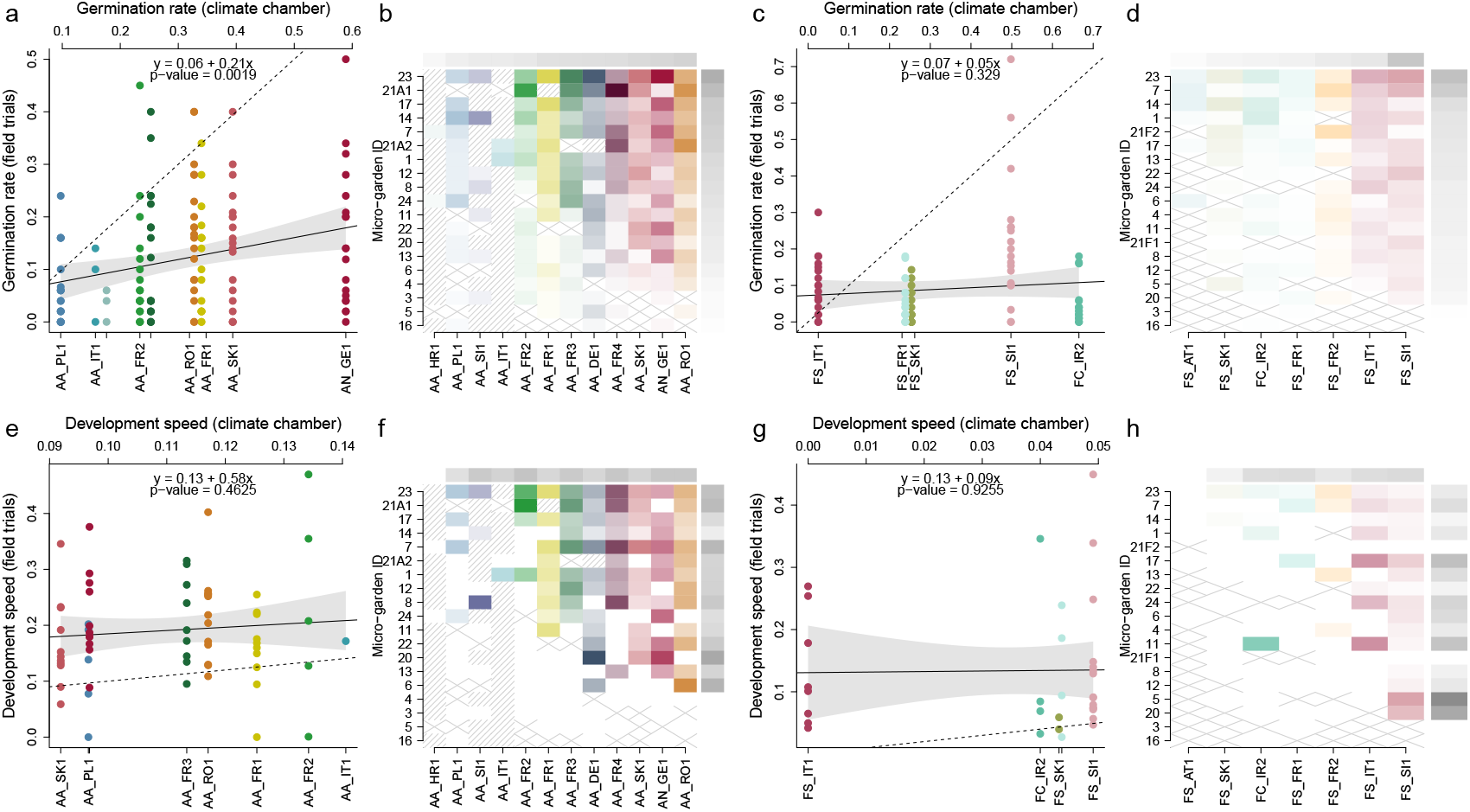
Germination rate (*g*) and development speed (*δ*) estimated from the field observations during the first growing season. Correlation between *g* and *δ* in the climate chambers and the field trials in fir (a) and (e) and in beech (c) and (g), respectively. The solid line indicates the estimated linear relationship between climate-chamber and field estimates, while the dashed lines show a one-to-one relationship. Provenance garden interaction in *g* and *δ* for fir (b) and (f) and beech (d) and (h), respectively. Color intensity corresponds to the value of *g* and *δ*, with darker colors indicating higher germination (*g*) and faster development (*δ*). Provenances and gardens are ordered by *g* and *δ*. The same provenance color code is used for fir and beech provenances. Marginals show the provenance or garden mean *g* and *δ*. Hatched cells indicate missing data, i.e., the provenance was not tested in the given garden. Cells with a cross indicate the lack of germination.

Comparisons between climate chamber and field performance revealed marked context dependence, with variation in germination rates among gardens often matching or exceeding that among provenances under controlled conditions (Fig. 3a, b). In fir, estimates of germination rate (*g*) from climate chambers and field trials were correlated but deviated substantially from a one-to-one relationship (Fig. 3a). On average, field germination reached only 21% of climate-chamber values, yet one-third of fir provenances exceeded their climate-chamber performance in at least one garden (e.g., AA PL1). In beech, no correspondence was detected between the climate chamber and field germination (Fig. 3b), and most provenances had sharply reduced germination in micro-gardens, including the provenance performing best in the climate chamber (FC IR2, 60%). But as in fir, some provenances exceeded their climate-chamber performance in individual gardens. This included the provenance FS IT1 that had very low germination under controlled conditions (*g <* 0.01) and was excluded from the survival analysis, but germinated at 30% in micro-garden 23.

Seedling development also differed markedly between environments. By the end of the first growing season, a larger fraction of seedlings in gardens had reached advanced developmental stages than in the climate chambers, likely reflecting longer exposure and higher accumulated growing degree days (Fig. S12b). Although average development was faster under controlled conditions, the range of development speeds (*δ* values) was substantially wider in the micro-gardens, consistent with strong environmental modulation of early growth trajectories (Fig. 3c,g).

### Environmental drivers of germination and first year survival

LASSO regression identified distinct sets of environmental predictors for maximum germination in the two genera. In fir, highly ranked predictors were primarily associated with micro-garden location, particularly soil properties. In beech, predictors reflected both provenance climate and micro-garden-level environmental conditions (Fig. S4). Mixed-effects and Markov models supported these patterns, indicating that gene–environment interactions explained the largest share of germination variability (Fig. 4a, Fig. 3b,d). In fir, most variation in germination was attributable to garden effects, especially fine-scale micro-environmental variation at the block level, whereas provenance effects were negligible (Fig. 4b). In beech, provenance explained a substantially larger share of variation, and environmental effects manifested primarily through interactions with provenance (Fig. 4c).

**Figure 4.**
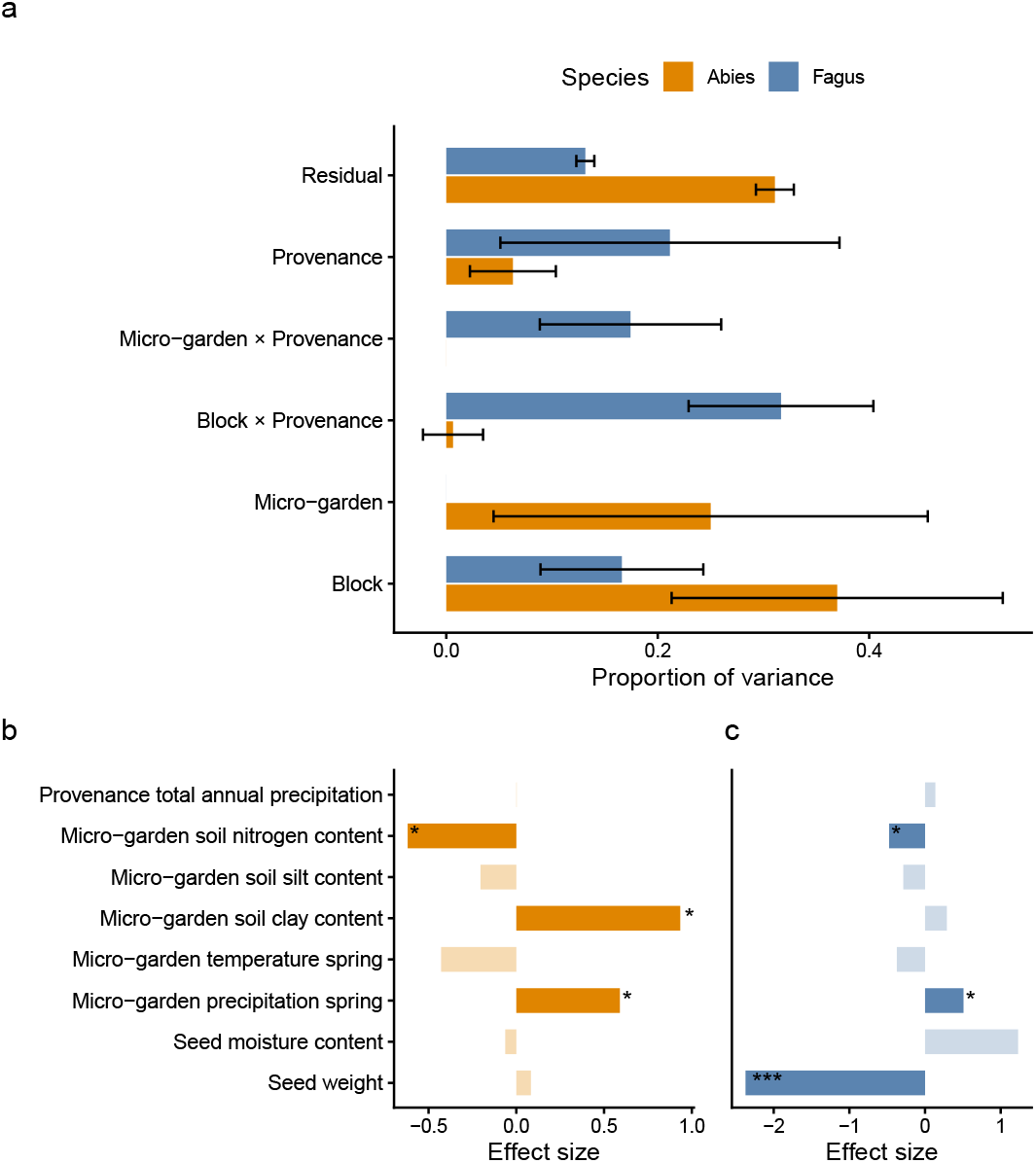
Variance partitioning and environmental drivers of maximum seed germination from the establishment of the experiment until 30 June. (a) Proportion of variance in germination explained by random effects for fir species and provenances (*Abies*, orange) and beech (*Fagus*, blue), including provenance, garden, block, their interactions, and residual variance. Error bars indicate the standard error of variance components. (b–c) Standardized effect sizes of fixed effects retained in the final models based on lasso regression for fir (b) and beech (c). Bars represent estimated coefficients from negative binomial mixed models; shading indicates non-significant effects. Asterisks denote significance based on Wald tests (*: p *<* 0.05, **: p *<* 0.01, ***: p *<* 0.001). Fixed effects are ordered by seed traits, garden climate, garden soil, and provenance climate variables.

Across both genera, spring precipitation positively influenced germination, while higher spring temperatures tended to reduce it even if the latter was not significant in the Wald test (Fig. 4b, c). Soil properties played an important role: nitrogen content negatively affected germination in both genera, and clay content had a strong positive effect in fir. In beech, in agreement with the climate chamber trials, germination was again strongly associated with seed weight, with lighter seeds germinating more successfully (Fig. 4b, c).

### Three-year survival of seedlings after germination

Seedling survival declined strongly with time since the cohort peak abundance in both fir and beech, with a highly significant effect of seasonal age bins (fir: Wald *χ*^2^ = 1107.3, *df* = 10, *P <* 0.001; beech: Wald *χ*^2^ = 631.7, *df* = 10, *P <* 0.001; Fig. 5a). Mortality was concentrated during the first summer following establishment and during subsequent winter and spring transition periods, after which survival trajectories flattened. As defined in the Methods, cohorts were tracked from the first census at which maximum post-germination abundance (*N*_0_) was observed, ensuring that temporal patterns reflect post-establishment survival rather than variation in germination timing. A modest increase in apparent survival during the second growing season (Fig. 5a) likely reflects a combination of detection error when seedlings present at one census are not observed at a previous census and the aggregation of observations within seasonal age bins.

**Figure 5.**
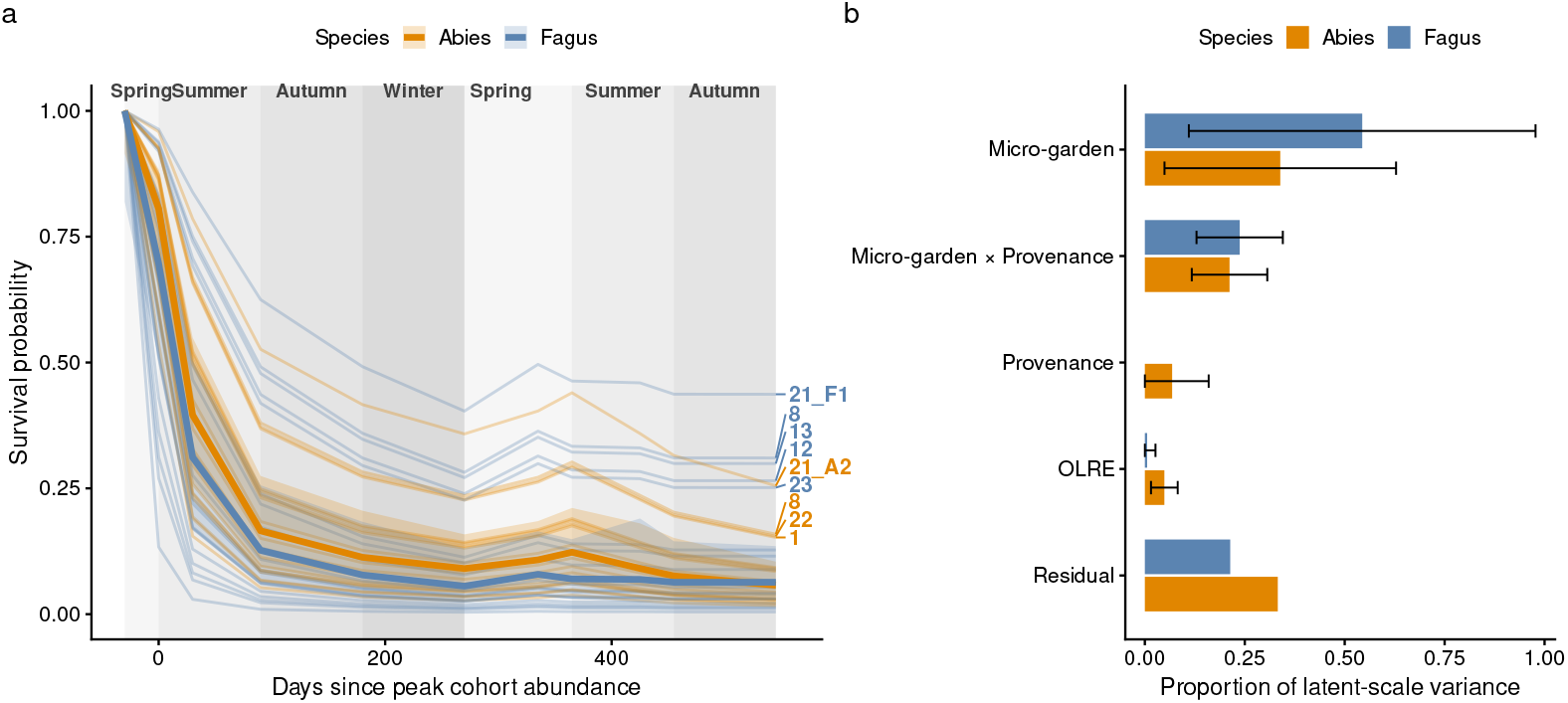
Modelled seedling survival of fir (*Abies*) and beech (*Fagus*) seedlings as a function of days since the peak abundance (*N*_0_) of the specific garden-provenance combination (cohort). Thin lines show garden-specific survival trajectories, while thick lines and shaded bands indicate genus-level mean survival and 95% confidence intervals. Background shading denotes seasonal periods relative to cohort peak. Garden identifiers are shown for gardens with predicted survival*≥* 0.15 at the final census.

Despite broadly similar temporal patterns of survival decline, fir and beech differed in the relative importance of micro-garden- and provenance-level effects, indicating genus-specific sensitivity to local environmental filters. Notably, micro-gardens supporting the highest late-stage survival were not consistently shared between genera: only gardens 21 and 8 showed high survival for both fir and beech (Fig. 5a), suggesting that distinct abiotic or biotic constraints shape post-establishment survival in the two genera.

Variance partitioning revealed substantial among-micro-garden variation in survival for both genera (Fig. 5b). Micro-garden effects and micro-garden–provenance interactions explained most of the variability, whereas provenance alone contributed comparatively little, and observation-level random effects were only detectable in fir. This pattern contrasts sharply with germination, where variance components were more evenly distributed among genetic and environmental sources and where provenance explained a substantial fraction of variation in beech (Fig. 4a). Together, these results indicate a clear shift in the dominant sources of variation across early life stages: germination is more strongly structured by genetic origin and fine-scale gene–environment interactions, whereas post-establishment survival is increasingly governed by broader, site-level environmental filters.

## Discussion

Seedling emergence and early establishment constitute major demographic bottlenecks in tree species across several biomes [Canham and Murphy, 2016, Rother et al., 2013]. While differential regeneration among species is well documented [Pacala et al., 1994, Bigelow and Canham, 2002], the processes structuring regeneration within species — particularly during the earliest life stages — remain poorly understood under natural forest conditions. This gap is striking given the long tradition of provenance and common-garden trials in forest trees [Savolainen et al., 2007, Alberto et al., 2013], which have generated extensive knowledge on genetic differentiation in growth and phenology, but typically bypass germination and early establishment by relying on nursery-grown seedlings. As a consequence, the strongest selective filters, which act at the start of the life cycle, are rarely observed directly [Muffler et al., 2021, de los Ángeles García-Hernández et al., 2019, Moser et al., 2025]. Our citizen-science-based experimental design, combined with controlled experiments, allowed us to follow cohorts continuously from seed to established seedling and to disentangle how genetic origin, local environmental conditions, and their interaction structure successive demographic filters. Our results reveal a consistent shift in the dominant drivers of regeneration across early life stages: while germination and early development are structured primarily by fine-scale gene–environment interactions, post-establishment survival is increasingly governed by broader site-level environmental filters (Fig. 3, 4, 5). Yet, fir and beech exhibited contrasting pathways through these early filters, reflecting differences in their broad life-history strategies and constraints.

Leites and Benito Garzón [2023] have emphasized the central role of gene–environment interactions (*G × E*) in forest tree provenance trials, arising through local adaptation and phenotypic plasticity. In line with this view, our results show that *G × E* already shape early life stages, influencing germination and early survival (Figs 3, 4, 5), but that their expression differs markedly between fir and beech. In fir, provenance differences in germination were relatively consistent across environments, and responses to environmental variation were broadly parallel among provenances. All fir provenances showed reduced germination under drought in the climate chamber (Fig. 2b), germination rates were correlated between climate chamber and field conditions (Fig. 3a). Although germination rate and development speed varied widely among field gardens (Fig. 3b,f), these differences were not associated with significant provenance-by-environment interactions (Fig. 4a), suggesting that micro-environmental heterogeneity, only partly captured by block effects, dominated early outcomes. A transplant experiment in silver fir provenances also found a prominent role of the local environment [Latreille and Pichot, 2017]. In contrast, we found that *G × E* became important at the survival stage, where garden-provenance interactions contributed substantially to variance after establishment (Fig. 5b).

In beech, by contrast, *G×E* effects were already prominent during germination and remained important through early survival. Beech provenances differed in their responses to drought in the climate chamber (Fig. 2e,f), and climate-chamber performance was a poor predictor of field germination (Fig. 3c,g), with several provenances germinating successfully only in specific gardens (Fig. 3e), which echos the findings of similar field trials [Muffler et al., 2021, Arend et al., 2016], and also consistent with high phenotypic plasticity in beech for physiological traits [Meier and Leuschner, 2008] and genomic evidence [Lazic et al., 2024]. Although germination generally increased with spring precipitation, provenance-related seed weight explained more variation than garden effects alone (Fig. 4c). Accordingly, block × provenance and garden × provenance interactions accounted for the largest share of variance in beech germination success (Fig. 4a), indicating strong context dependence already at the earliest life stage.

Consistent with this framework, germination timing showed a clear geographic structure in both genera, with longitude significantly affecting germination hazard under controlled conditions (Fig. 2c,f). Geographic predictors likely integrate broad-scale variation in dormancy, chilling requirements, and thermal thresholds, traits commonly shaped by adaptation to local climatic conditions [Bevington, 1986, Donohue et al., 2010, Walck et al., 2011]. While geographic effects on germination timing were consistently expressed in climate chambers, their translation to the field depended on genus-specific regeneration strategies. Seed mass emerged as a key modulator of this translation: as a key functional trait, it reflects fundamental trade-offs between fecundity and dispersal [Westoby et al., 1996] and determines early developmental trajectories [Donohue et al., 2010]. In fir, seed weight covaried with longitude (Fig. S7a), suggesting broader geographic differentiation between populations owing to their colonization history [Pistone et al., 2025], during which multiple correlated traits related to seed development and dormancy could have co-evolved. In beech, by contrast, seed weight was independent of longitude and acted as the most important factor shaping germination responses across environments (Fig. S7b; Fig. 4c). Moreover, seed weight influenced early post-germination development in both species: lighter seeds developed faster, reflected by higher thermal time required to progress through germination stages (*δ*) under controlled conditions (Fig. S7). This suggests that seed mass not only affects the probability of germination, but also modulates the rate at which seedlings traverse the vulnerable early life stages, potentially related to colonization dynamics [Coomes and Grubb, 2003]. Indeed, in an intraspecific context, small-seeded species favor rapid emergence and are better at colonizing, while large-seeded species rely more on successful establishment and perform better in a competitive environment [Clark et al., 2004, Turnbull et al., 1999, Qiu et al., 2021].

Global syntheses indicate that germination timing is often more responsive to temperature and climatic gradients than final germination percentage [Vicente and Benito Garzón, 2024]. Consistent with this pattern, we found that germination speed (*t*_50_), synchrony, and phenological development speed (*δ*), were more responsive to environmental variation and provenance interactions than cumulative germination. Accordingly, development speed estimated under climate-chamber conditions was a poor predictor of development in the forest, where development was also considerably slower (Fig. 3c,g), indicating that early germination phenology integrates fine-scale environmental cues that are homogenized or absent under controlled conditions. A similar decoupling between regional climate signals and local phenological expression has been demonstrated for budburst phenology, in which microclimate and resource availability strongly modulate responses to temperature [Vitasse et al., 2021]. Together, these patterns reinforce the role of phenology as a buffer against temporal environmental heterogeneity [Chuine, 2010], and agree with findings in silver fir, where provenances’ budburst phenological trajectories were decoupled from growth and drought tolerance in seedlings [Csilléry et al., 2020a].

Regeneration in trees is highly sensitive to drought, soil moisture, and microsite conditions [Koelemeijer et al., 2024, Phillips et al., 2016, De Frenne et al., 2013, Valladares et al., 2016, Zellweger et al., 2020], resulting in characteristically patchy recruitment patterns that reflect fine-scale heterogeneity in microclimate, soil properties, and biotic interactions [Clark et al., 1999]. Such spatial heterogeneity creates distinct regeneration niches [Beckage and Clark, 2003, Atkins et al., 2023, Mwamulima et al., 2025], and interactions between canopy structure and understory environments are well known to shape seedling performance, species composition, and competitive dynamics, including under drought stress [Pacala et al., 1994, Bigelow and Canham, 2002, Walters et al., 2023]. Consistent with this framework, our results show that micro-environmental variation was a dominant driver of germination under natural conditions: block-level effects ( 25 m^2^) explained a substantial proportion of variation in germination success (Fig. 4a). In beech, this fine-scale heterogeneity further interacted with provenance-specific seed size (Fig. 4c), highlighting how local environmental variation can modulate trait effects at the earliest life stages.

At the same time, we detected a consistent negative association between germination success and soil nitrogen content derived from SoilGrids (Fig. 4b,c), indicating that broader edaphic context can also constrain regeneration. Because these nitrogen estimates represent landscape-scale soil fertility rather than microsite conditions [de Sousa et al., 2021], this pattern likely reflects integrative site-level properties of productive forest soils, such as dense understory vegetation or elevated pathogen pressure [Aber, 1992], which can reduce germination success despite being favorable for later growth stages. As seedlings grow and rooting depth and exposure increase, the relative influence of microsite conditions and fine-scale gene–environment interactions is expected to decline, while broader site-level environmental filters become increasingly important [Chang-Yang et al., 2021]. In line with this expectation, we observed a clear shift in the dominant sources of variation across early life stages: germination was structured primarily by fine-scale environmental heterogeneity and gene–environment interactions, whereas post-establishment survival was increasingly governed by site-level conditions (Fig. 5), consistent with the regeneration niche concept [Grubb, 1977].

Consistent with the shift from fine-scale germination filters to broader environmental control during establishment, seedling survival declined steeply during the first summer after establishment (Fig. 5a), likely due to heat- and drought-related stress. Additional losses occurred during winter and spring thaw, when freeze injury and frost heave are possible. A slight increase in apparent survival in early summer of the second year likely reflects imperfect detectability and/or changing cohort representation among later censuses rather than true recruitment (Fig. 5a). To summarize our findings in numbers, we found that of the 18,000 seeds sown across the experiment, fewer than 10% (n = 1,716) germinated. This selection filter depends on fecundity, masting, and, in our case, potentially also seed lot effects related to drying and storage. The second major filter occurred during the first spring and summer, when again fewer than 10% of the germinated seedlings survived (n = 167) until the end of the first vegetative season. Thereafter, survival probability curves became comparatively flat (Fig. 5a). By quantifying survival probabilities across a heterogeneous range of environments, our experimental approach improves understanding of selection filters across life stages, overcomes sampling limitations inherent to observational studies of natural regeneration [Clark et al., 1999, Johnson et al., 2008], and provides empirical inputs for demographic and forest dynamics models [Díaz-Yañez et al., 2024].

Successful regeneration underpins future forest composition and dynamics, and variation in recruitment plays a central role in shaping forest responses to climate change [Anderson-Teixeira et al., 2013]. While macroclimate warming can be partially buffered by forest canopies [De Frenne et al., 2013, Zellweger et al., 2020], our results show that early regeneration is already filtered by local environments in ways that interact with provenance-level genetic variation. Although within-genus provenance effects on germination were comparatively small, their persistence indicates that genetic variation expressed during regeneration can influence the future genetic composition of forests. This finding has direct implications for assisted migration programs, which commonly rely on nursery-grown seedlings for both planning and implementation, thereby bypassing the strongest early selective filters [Chakraborty et al., 2024, Hällfors et al., 2014, Mauri et al., 2023]. Given increasing reports of climate-induced regeneration failure across Europe and beyond [Talluto et al., 2017, Rumpf et al., 2019], incorporating germination and early establishment into adaptation planning will be essential to improve the ecological realism and long-term success of assisted migration strategies and may foster regeneration-based management such as direct seeding [Ceccon et al., 2016, Grossnickle and Ivetić, 2017].

## Acknowledgments

This project was funded by the ERC Consolidator Grant MyGardenOfTrees (grant agreement no. 101003296) awarded to K.C., which supported the work of J.C.d.S., M.Ce., N.C., and M.B., and contributed to the work of M.Ca. We are deeply grateful to the voluntary foresters who established micro-gardens and carried out the field observations. We thank Gabor Reiss for valuable advice on the climate chamber setup. We also thank Haonan Yang for his contributions to the development of observation forms using ODK Collect, and Christoph Sperisen and Yann Vitasse for insightful discussions during the planning of the experiments and for input that helped advance the interpretation of the results.

## Author Contributions

K.C. conceived the study, coordinated the project, designed the experimental protocols, obtained funding, performed the final data analyses, integrated all analytical components, produced the figures, and wrote the manuscript. J.C.d.S. collected field and environmental data, curated and validated the datasets, and contributed to data processing and analyses. M.Ca. developed the Markov model, estimated its parameters, and contributed to visualization. J.A. conducted climate chamber experiments, contributed to data processing and analyses, and provided input through his MSc thesis work. M.B. collected field data and contributed to data processing and analyses. M.Ce. conducted climate chamber experiments and contributed to data processing and analyses. N.P. led the development of the experimental protocols, coordinated the purchase of seeds and experimental materials, coordinated the climate chamber and field experiments, collected field data, and supervised data collection by foresters. D.W. developed and supervised the Markov model, contributed to the interpretation of the results, and provided input on the manuscript. All authors reviewed and approved the final manuscript.

## Data availability

All data and code supporting the findings of this study are openly available. Data analysis scripts and processed datasets from the *MyGardenOfTrees* pilot trials (climate chamber and micro-garden experiments) are hosted in a public GitHub repository at https://github.com/kcsillery/MyGardenOfTrees_PilotTrials/tree/main.

The model and inference scheme were implemented as a command-line C++ program, tree growth, using the stattools library. The source code and user manual are available at https://bitbucket.org/wegmannlab/tree_growth/. All analyses reported here were conducted using commit a9ae485.

## Supporting Information

Additional Supporting Information may be found in the online version of this article.

| File | Description |
| --- | --- |
| Figures_S1-14_Table_S1.pdf | Fig. S1 Climate chamber trays for fir (*Abies*) and beech (*Fagus*) germination. |
|  | Fig. S2 Phenological stages used to score germination and early seedling development. |
|  | Fig. S3 LASSO-selected environmental predictors of germination in fir and beech. |
|  | Fig. S4 Standardized regression coefficients of LASSO-selected predictors. |
|  | Fig. S5 Climate chamber germination trajectories across provenances. |
|  | Fig. S6 Relationships between seed density and germination metrics. |
|  | Fig. S7 Correlations among germination metrics, geography, and seed traits. |
|  | Fig. S8 Effect of experimental cycle length on germination metrics. |
|  | Fig. S9 Cumulative germination as a function of growing degree-days in micro-gardens. |
|  | Fig. S10 Field germination trajectories by provenance and micro-garden. |
|  | Fig. S11 Fir (*Abies*) phenological development in climate chambers and gardens. |
|  | Fig. S12 Beech (*Fagus*) phenological development in climate chambers and gardens. |
|  | Fig. S13 Distribution of growing degree-days across gardens and provenances. |
|  | Fig. S14 Diagnostic plots for mixed-effects models of germination. |
|  | Table S1 Germination metrics across species, provenances, treatments, and experimental cycles. |
| Methods_S1.pdf | Protocols for establishing and monitoring the *MyGardenOfTrees* climate chamber and micro-garden experiments. |
| Methods_S2.pdf | Description of a hidden Markov model of seed germination, phenological development, and mortality, including the inference and importance sampling scheme. |
| Dataset_S1.zip | XLSForms and associated media files used for standardized data collection in the *MyGardenOfTrees* citizen science project. |

