## Supplementary material for "Gene–environment interactions govern early regeneration in fir and beech: evidence from participatory provenance trials across Europe": Methods S2

### Description of a hidden Markov model of seed germination, phenological development, and mortality, including the inference and importance sampling scheme

Let $z_i(t)$ denote the life stage of seed $i$ on day $t$. We consider three life stages: i) a viable seed that is dormant and has not yet germinated ($z_i(t) = \mathcal{L}_\mathcal{S}$), ii) a seed that germinated into a currently growing seedling ($z_i(t) = \mathcal{L}_\mathcal{G}$) and iii) non-viable seed or a previously germinated seedling that has died ($z_i(t) = \mathcal{L}_\mathcal{D}$). On each day, four events may happen in this order: 1) a seedling will grow, 2) a seedling may die, 3) a seed may germinate and 4) a seed or seedling may be observed. We denote by $t_i^{(g)}$ and $t_i^{(d)}$ the days on which seedling $i$ germinated and died, respectively.

We assume all seedlings can be partitioned into $K$ classes (e.g. combination of provenance and growing location) and let $c_i \in \mathcal{C} = \{\mathcal{C}_1, \ldots, \mathcal{C}_K\}$ denote the class of seedling $i$.

##### Germination

Let $g_c$ be the probability that a seed of class $c \in \mathcal{C}$ is viable and can germinate. Thus, $z_i(0) = \mathcal{L}_\mathcal{S}$ with probability $g_{c_i}$ and $z_i(0) = \mathcal{L}_\mathcal{D}$ with probability $1 - g_{c_i}$.

We assume the probability that a seed germinates, i.e. that it transitions from $z_i(t-1) = \mathcal{L}_\mathcal{S}$ to $z_i(t) = \mathcal{L}_\mathcal{G}$, to depend on the growing degree days $h(t)$ up to day $t$. We model this dependence using the logistic function

$$\mathbb{P}(z_i(t) = \mathcal{L}_\mathcal{G} | z_i(t-1) = \mathcal{L}_\mathcal{S}, \boldsymbol{G}_{c_i}) = \frac{1}{1 + \exp[-\gamma_{c_i}(h(t) - \bar{h}_{c_i})]},$$

where $\boldsymbol{G}_{c_i} = (g_{c_i}, \gamma_{c_i}, \bar{h}_{c_i})$ denotes the vector of parameters relevant for germination, of which the logistic growth rate $\gamma_{c_i}$ determines the variation in germination among seeds of a class and $\bar{h}_{c_i}$ the growing degree days necessary to reach a germination probability of $\frac{1}{2}$.

The probability that seed $i$ germinated on day $t_i^{(g)}$ is given by

$$\begin{aligned} \mathbb{P}(t_i^{(g)}|\boldsymbol{G}_{c_i}) &= g_{c_i}\mathbb{P}(z_i(t_i^{(g)}) \\ &= \mathcal{L}_{\mathcal{G}}|z_i(t_i^{(g)}-1) = \mathcal{L}_{\mathcal{S}}, \boldsymbol{G}_{c_i}) \prod_{t=1}^{t_i^{(g)}-1} \mathbb{P}(z_i(t) = \mathcal{L}_{\mathcal{S}}|z_i(t-1) = \mathcal{L}_{\mathcal{S}}, \boldsymbol{G}_{c_i}), \end{aligned} \tag{1}$$

where

$$\mathbb{P}(z_i(t) = \mathcal{L}_{\mathcal{S}}|z_i(t-1) = \mathcal{L}_{\mathcal{S}}, \boldsymbol{G}_{c_i}) = 1 - \mathbb{P}(z_i(t) = \mathcal{L}_{\mathcal{G}}|z_i(t-1) = \mathcal{L}_{\mathcal{S}}, \boldsymbol{G}_{c_i}).$$

#### Development

Let $x_i(t)$ quantify the size of a currently growing $(z_i(t) = \mathcal{L}_{\mathcal{G}})$ seedling $i$ on day $t$. We assume that newly germinated seedlings have a size of zero and grow linearly after germination with class-specific growth rate $\delta_c$ such that for a seed $i$ with class $c_i$

$$x_i(t) = \begin{cases} \delta_{c_i}(t - t_i^{(g)}) & \text{if } z_i(t) = \mathcal{L}_{\mathcal{G}} \text{ and } t_i^{(g)} < t \leq t_i^{(d)} \\ 0 & \text{otherwise} \end{cases}. \tag{2}$$

Note that the variable $x_i(t)$, while referred to as "size", does not mean an actual measurable size (e.g. tree height), but is used here just as a measure of how a seedling is moving along development stages. Note also that while modeled linearly, we allow for a non-linear translation to life stages (see below).

#### Death

A currently growing seedling $i$ dies with a probability that is dependent on its current life stage. We assume the highest death rate occurs at germination and then declines exponentially with size. For a seedling $i$ of class $c_i$ we thus have

$$\mathbb{P}(z_i(t) = \mathcal{L}_{\mathcal{D}}|z_i(t-1) = \mathcal{L}_{\mathcal{G}}, \boldsymbol{D}_{c_i}) = \alpha_{c_i} \exp[-\beta_{c_i} x_i(t)],$$

where $\boldsymbol{D}_{c_i} = (\alpha_{c_i}, \beta_{c_i})$ denotes the vector of parameters relevant for death and

$$\mathbb{P}(z_i(t) = \mathcal{L}_{\mathcal{G}}|z_i(t-1) = \mathcal{L}_{\mathcal{G}}, \boldsymbol{D}_{c_i}) = 1 - \mathbb{P}(z_i(t) = \mathcal{L}_{\mathcal{D}}|z_i(t-1) = \mathcal{L}_{\mathcal{G}}, \boldsymbol{D}_{c_i}).$$

The probability that seed $i$ died on day $t_i^{(d)}$ is given by

$$\mathbb{P}(t_i^{(d)}|t_i^{(g)}, \boldsymbol{D}_{c_i}) = \prod_{t=t_i^{(g)}+1}^{t_i^{(d)}-1} \Big[\mathbb{P}(z_i(t) = \mathcal{L}_\mathcal{G}|z_i(t-1) = \mathcal{L}_\mathcal{G}, \boldsymbol{D}_{c_i})\Big] \mathbb{P}(z_i(t_i^{(d)}) = \mathcal{L}_\mathcal{D}|z_i(t_i^{(d)}-1) = \mathcal{L}_\mathcal{G}, \boldsymbol{D}_{c_i}). \tag{3}$$

Note that since $x_i(t) = x_i(t-1) + \delta_{c_i}$, these probabilities can be calculated iteratively for efficiency:

$$\begin{aligned} \exp[-\beta_{c_i} x_i(t)] &= \exp[-\beta_{c_i}((x_i(t-1) + \delta_{c_i})] \\ &= \exp[-\beta_{c_i} x_i(t-1)] \exp[-\beta_{c_i}\delta_{c_i}]. \end{aligned}$$

This iteration only involves the constant term $\exp[-\beta_{c_i}\delta_{c_i}]$, and no exponentials need to be evaluated.

#### Emission probabilities: the observed life stages

Let $\tau_1, \ldots, \tau_M$ denote $M$ times at which seeds were measured, and let $\tau_0 < \tau_1$ denote the beginning of the experiment. Let $\boldsymbol{s}_i = (s_i(\tau_1), \ldots, s_i(\tau_M))$ denote the life stage identified for seedling $i$ identified at the observation times with $s_i(\tau_m) \in \mathcal{S} = \{\mathcal{S}_0, \mathcal{S}_1, \ldots, \mathcal{S}_S\}$. Of these, $\mathcal{S}_0$ denotes the unobserved state, i.e. the seedling has not yet germinated or has already died. For all other stages $\mathcal{S}_s, s > 0$, we model emission probabilities as normalized gamma densities. Let $w_s(x) = \text{Gamma}(x; m_s, \sigma_s^2)$ be the weight of stage $s$ for seedling size $x$, where $\boldsymbol{m} = (m_1, \ldots, m_S)$ and $\boldsymbol{\sigma}^2 = (\sigma_1^2, \ldots, \sigma_S^2)$ are the modes and variances of the gamma distribution relevant for stages $s = 1, \ldots, S$. Denoting by $\boldsymbol{E} = (\boldsymbol{m}, \boldsymbol{\sigma}^2)$ the vector of emission parameters, the emission probabilities are then given by

$$\mathbb{P}(s_i(\tau_m) = \mathcal{S}_s|z_i(\tau_m), x_i(\tau_m), \boldsymbol{E}) = \begin{cases} 1 & \text{if } z_i(\tau_m) = \mathcal{L}_\mathcal{S} \text{ and } s = 0 \\ 1 & \text{if } z_i(\tau_m) = \mathcal{L}_\mathcal{D} \text{ and } s = 0 \\ \frac{w_s(x_i(\tau_m))}{\sum_l w_l(x_i(\tau_m))} & \text{if } z_i(\tau_m) = \mathcal{L}_\mathcal{G} \\ 0 & \text{otherwise} \end{cases} \tag{4}$$

We assume that the stage $s_\mathcal{D} \in \mathcal{S}$ with the highest mortality is known and set $m_{\mathcal{S}_S} = x_\mathcal{D}$. Note that the model presented is not identifiable, as multiple combinations of modes $m_s$ and growth rates $\delta_c$ can lead to the same emissions. To avoid these non-identifiability issues, we will arbitrarily assume $m_{\mathcal{S}_S} = x_\mathcal{D} = 100$.

The emission probability of the full vector of observations is given by

$$\mathbb{P}(\boldsymbol{s}_i|t_i^{(g)}, \boldsymbol{z}_i, \delta_{c_i}, \boldsymbol{E}) = \prod_{m=1}^{M} \mathbb{P}(s_i(\tau_m)|z_i(\tau_m), \delta_{c_i}(\tau_m - t_i^{(g)}), \boldsymbol{E})$$

### Probability of full realizations

Given the above model, the seeds follow a Markov model with class-specific parameters $\boldsymbol{\theta}_c = (\boldsymbol{G}_c, \delta_c, \boldsymbol{D}_c)$ as well as the emission parameters $\boldsymbol{E}$. For seed $i$, we distinguish three different types of realizations $\boldsymbol{z}_i = (z_i(0), \ldots, z_i(\tau_M))$ until time $\tau_M$:

1. The seed may be viable, germinate between time points $m-1$ and $m$ and be observed for the remainder of the experiment at time points $\tau_m, \ldots, \tau_M$. We obtain the probability of the observations by integrating out the specific day of germination $\tau_{m-1} \leq t_i^{(g)} < \tau_m$:

$$\mathbb{P}\Big(\boldsymbol{s}_i, z_i(\tau_0, \ldots, \tau_{m-1}) = \mathcal{L}_{\mathcal{S}}, z_i(\tau_m, \ldots, \tau_M) = \mathcal{L}_{\mathcal{G}}|\boldsymbol{\theta}_{c_i}, \boldsymbol{E}\Big) = \\ \sum_{t^{(g)}=\tau_{m-1}}^{\tau_m} \mathbb{P}(t^{(g)}|\boldsymbol{G}_{c_i}) \prod_{t=t^{(g)}+1}^{\tau_M} \Big[\mathbb{P}(z_i(t) = \mathcal{L}_{\mathcal{G}}|z_i(t-1) = \mathcal{L}_{\mathcal{G}}, \boldsymbol{D}_{c_i})\Big] \mathbb{P}(\boldsymbol{s}_i|t^{(g)}, \boldsymbol{z}_i, \boldsymbol{E}, \delta_{c_i}),$$

   where we used the notation $z_i(t_1, t_2, \ldots) = \mathcal{L}$ to indicate that the life stage was $\mathcal{L}$ at all times $t_1, t_2, \ldots$ specified.

2. The seed may be viable, germinate between time points $m-1$ and $m$, be observed at time points $\tau_m, \ldots, \tau_n$ and die prior to time point $\tau_{n+1} <= \tau_M$. We obtain the probability of the observations by integrating out the specific days of germination $\tau_{m-1} \leq t_i^{(g)} < \tau_m$ and death $\tau_n < t_i^{(d)} \leq \tau_{n+1}$:

$$\mathbb{P}\Big(\boldsymbol{s}_i, z_i(\tau_0, \ldots, \tau_{m-1}) = \mathcal{L}_{\mathcal{S}}, z_i(\tau_m, \ldots, \tau_n) = \mathcal{L}_{\mathcal{G}}, z_i(\tau_{n+1}, \ldots, \tau_M) = \mathcal{L}_{\mathcal{D}}|\boldsymbol{\theta}_{c_i}, \boldsymbol{E}\Big) = \\ \sum_{t^{(g)}=\tau_{m-1}}^{\tau_m} \mathbb{P}(t^{(g)}|\boldsymbol{G}_{c_i}) \mathbb{P}(\boldsymbol{s}_i|t^{(g)}, \boldsymbol{z}_i, \boldsymbol{E}, \delta_{c_i}) \sum_{t^{(d)}=\tau_n+1}^{\tau_{n+1}} \mathbb{P}(t^{(d)}|t^{(g)}, \boldsymbol{D}_{c_i})$$

3. The seed was never observed at any of the time points $\tau_1, \ldots, \tau_M$. This may occur due to three distinct reasons:

   (a) The seed may be non-viable and stay in state $z_i(t) = \mathcal{L}_{\mathcal{D}}$ for all $t = 0, \ldots, \tau_M$. This occurs with probability

$$\mathbb{P}\Big(s_i(\tau_1, \ldots, \tau_M) = \mathcal{S}_0, z_i(\tau_1, \ldots, \tau_M) = \mathcal{L}_{\mathcal{D}}|\boldsymbol{\theta}_{c_i}, \boldsymbol{E}\Big) = 1 - g_{c_i}$$

(b) The seed may be viable and never germinate. This occurs with probability

$$\mathbb{P}\Big(s_i(\tau_1,\dots,\tau_M)=\mathcal{S}_0, z_i(\tau_1,\dots,\tau_M)=\mathcal{L}_\mathcal{S}|\boldsymbol{\theta}_{c_i},\boldsymbol{E}\Big) = g_c \prod_{t=1}^{\tau_M} \mathbb{P}(z_i(t)=\mathcal{L}_\mathcal{S}|z_i(t-1)=\mathcal{L}_\mathcal{S},\boldsymbol{G}_{c_i})$$

(c) The seed may be viable, germinate between time points $m-1$ and $m$ and die before timepoint $m$. We obtain the probability of the observations by integrating out the specific days of germination $\tau_{m-1} \leq t_i^{(g)} < \tau_m - 1$ and death $t_i^{(g)} < t_i^{(d)} \leq \tau_m$:

$$\begin{aligned}&\mathbb{P}\Big(s_i(\tau_1,\dots,\tau_M)=\mathcal{S}_0, z_i(\tau_0,\dots,\tau_{m-1})=\mathcal{L}_\mathcal{S}, z_i(\tau_m,\dots,\tau_M)=\mathcal{L}_\mathcal{D}|\boldsymbol{\theta}_{c_i},\boldsymbol{E}\Big)\\ &= \sum_{t^{(g)}=\tau_{m-1}}^{\tau_m-1} \mathbb{P}(t^{(g)}|\boldsymbol{G}_{c_i}) \sum_{t^{(d)}=t^{(g)}+1}^{\tau_m} \mathbb{P}(t^{(d)}|t^{(g)},\boldsymbol{D}_{c_i}).\end{aligned}$$

The probability of $\mathbb{P}\Big(s_i(\tau_1,\dots,\tau_M)=\mathcal{S}_0)|\boldsymbol{\theta}_{c_i},\boldsymbol{E}\Big)$ is thus given by

$$\begin{aligned}&\mathbb{P}\Big(s_i(\tau_1,\dots,\tau_M)=\mathcal{S}_0)|\boldsymbol{\theta}_{c_i},\boldsymbol{E}\Big) =\\ &\mathbb{P}\Big(s_i(\tau_1,\dots,\tau_M)=\mathcal{S}_0, z_i(\tau_1,\dots,\tau_M)=\mathcal{L}_\mathcal{D}|\boldsymbol{\theta}_{c_i},\boldsymbol{E}\Big)\\ &+\mathbb{P}\Big(s_i(\tau_1,\dots,\tau_M)=\mathcal{S}_0, z_i(\tau_1,\dots,\tau_M)=\mathcal{L}_\mathcal{S}|\boldsymbol{\theta}_{c_i},\boldsymbol{E}\Big)\\ &+\sum_{m=1}^{M-1}\mathbb{P}\Big(s_i(\tau_1,\dots,\tau_M)=\mathcal{S}_0, z_i(\tau_0,\dots,\tau_{m-1})=\mathcal{L}_\mathcal{S}, z_i(\tau_m,\dots,\tau_M)=\mathcal{L}_\mathcal{D}|\boldsymbol{\theta}_{c_i},\boldsymbol{E}\Big)\end{aligned}$$

### Micro-gardens in which seeds were not individually tracked

In our micro-garden experiment, seeds were not individually tracked and hence only the counts of life-stages across a set of seedlings of a specific class grown together as one set are available. Let $\boldsymbol{n}_{j\tau_m} = (n_{j\tau_m 1},\dots,n_{j\tau_m S})$ denote the vector of stage counts for set $j$ of size $|\boldsymbol{n}_j|$ at time $\tau_m$ and let $\boldsymbol{n}_j = (\boldsymbol{n}_{j\tau_1},\dots,\boldsymbol{n}_{j\tau_M})$. Let us further denote by $\boldsymbol{S}$ the matrix of stage assignments to seeds in set $j$ with elements $S_{[im]}$ denotes the stage of seed $i$ at time $\tau_m$. Rows $\boldsymbol{s}_i$ of $\boldsymbol{S}$ thus correspond to the observations of seed $i$. To calculate the probability $\mathbb{P}(\boldsymbol{n}_j|\boldsymbol{\theta}_{c_i},\boldsymbol{E})$ of all observations of $\boldsymbol{n}_j$ of that set across all time points, we need to integrate over all assignments of stages to seeds $\boldsymbol{S}$:

$$\mathbb{P}(\boldsymbol{n}_j|\boldsymbol{\theta}_{c_i},\boldsymbol{E}) = \sum_{\boldsymbol{S}} \mathbb{P}(\boldsymbol{n}_j|\boldsymbol{S})\mathbb{P}(\boldsymbol{S}|\boldsymbol{\theta}_{c_i},\boldsymbol{E}),$$

where

$$\mathbb{P}(\boldsymbol{S}|\boldsymbol{\theta}_{c_i}, \boldsymbol{E}) = \prod_i \mathbb{P}(\boldsymbol{s}_i|\boldsymbol{\theta}_{c_i}, \boldsymbol{E})$$

and

$$\mathbb{P}(\boldsymbol{n}_j|\boldsymbol{S}) = \begin{cases} 1 & \text{if } n(\boldsymbol{S}) = \boldsymbol{n}_j \\ 0 & \text{otherwise} \end{cases},$$

that is, $\mathbb{P}(\boldsymbol{n}_j|\boldsymbol{S}) = 1$ if the stage counts per time point $n(\boldsymbol{S})$ match the observed counts $\boldsymbol{n}_j$ and zero otherwise.

The space of potential stage assignments $\boldsymbol{S}$ is very large even for a small number of seeds and cannot be explored in full. We therefore use a Markov approximation to integrate over stage assignments. Since a naive sampling of state assignments $\boldsymbol{S} \sim \mathbb{P}(\boldsymbol{S}|\boldsymbol{\theta}_{c_i}, \boldsymbol{E})$ results in a larger number of samples that are incompatible with the data, we generate samples of $\boldsymbol{S}$ using importance sampling:

$$\mathbb{P}(\boldsymbol{n}_j|\boldsymbol{\theta}_{c_i}, \boldsymbol{E}) \approx \sum_{k=1}^{K} w_k \mathbb{P}(\boldsymbol{n}_j|\boldsymbol{S}_k)$$
$$\boldsymbol{S}_k \sim \mathbb{P}_I(\boldsymbol{S}|\boldsymbol{\theta}_{c_i}, \boldsymbol{E})$$
$$w_k = \frac{\mathbb{P}(\boldsymbol{S}|\boldsymbol{\theta}_{c_i}, \boldsymbol{E})}{\mathbb{P}_I(\boldsymbol{S}|\boldsymbol{\theta}_{c_i}, \boldsymbol{E})}.$$

We simulate a fraction $\pi$ of samples under the full model, i.e. by simulating for each seed whether it germinated or not with probability $g_c$, and if it did germinate, the day of germination according to (1) and the day of death according to (3). These samples have weight $w_k = \frac{1}{\pi}$.

We generate the remaining fraction $1 - \pi$ of samples as follows:

- Let $\tilde{\boldsymbol{g}}_k$ denote a sorted vector of sampled days at which a seeds germinated. The same day may occur multiple times and if all seeds of a set germinated, its length matches that of the number if seeds of a specific class. Let $\tilde{\boldsymbol{d}}_k$ analogously be a sorted vector of the intervals (prior to the first, after the last or between consecutive observation times) during which death occurred. We fill these containers as follows: We first identify the minimal number of germination events per interval and pick a random germination day within that interval. If three seeds were observed at time observation time $m$ but four seeds at time $m + 1$, for instance, at least one germination event must have occurred between these observation times and we pick a random day according to (1), truncated to that interval:

$$\mathbb{P}(t_i^{(g)}|\boldsymbol{G}_{c_i}, \tau_m \le g < \tau_{m+1}) = \frac{\mathbb{P}(t_i^{(g)}|\boldsymbol{G}_{c_i})}{\sum_{g'=\tau_m}^{\tau_{m+1}-1} \mathbb{P}(t_i^{(g')}|\boldsymbol{G}_{c_i})}.$$

We analogously identify the minimal number of death events that occurred in each interval and add those intervals to $\tilde{\boldsymbol{d}}_k$.

While it is unknown if and when the remaining seeds germinated, the number of additional germination and death events must be equal in each interval. For each of the unaccounted seeds we therefore i) first simulate if the seed is germinating or not with probability $g_c$, ii) then, in case the seed germinated, simulate a random germination day (without any interval restriction) according to (1) and add a death event to the interval in which the germination was simulated.

- We next simulate sample pairs of germination dates and death intervals given the constraint that germination must predate death. We start with the earliest death interval $\tilde{d}_{k1}$ of $\tilde{\boldsymbol{d}}_k$ (the first since the vector is sorted) and randomly pick a germination date from $\tilde{\boldsymbol{g}}_k$ that is compatible with a death in that interval, i.e. from $\tilde{\boldsymbol{g}}'_k = (\tilde{g}_{ki} | \tilde{g}_{ki} < \tau_{\tilde{d}_{k1}})$. Once the germination data was simulated, we simulate a valid death date in the chosen death interval that postdates the simulated germination date. We repeat that process for each death interval in sequence.
- We calculate the importance weights $w_k$ as $\frac{1}{1-\pi}$ times the ration between the probability of the simulated $\boldsymbol{S}_k$ under the full model and its probability under the importance sampling scheme outlined above.

In each iteration, we initially generate $20,000$ samples and determine the effective sample size ESS of the samples generated as

$$ESS = \frac{(\sum_k w_k)^2}{\sum_k w_k^2}.$$

We then generate an additional batches of $20,000$ samples until the ESS of all samples combined exceeds $2,000$.

### Implementation

We implemented the above model and inference scheme as a command-line `C++` program `tree_growth` using the library `stattools`. The implementation, along with a brief user manual, is available at `https://bitbucket.org/wegmannlab/tree_growth/`. All estimations in this paper were done with commit `a9ae485`.
