## Supplementary material for "Gene–environment interactions govern early regeneration in fir and beech: evidence from participatory provenance trials across Europe": Methods S1

Swiss Federal Institute for Forest, Snow and Landscape Research

European Commission

European Research Council
Established by the European Commission

### MyGardenOfTrees

#### A participatory science project for climate-resilient forests in Europe

##### Trials 2021-2026

#### Protocol

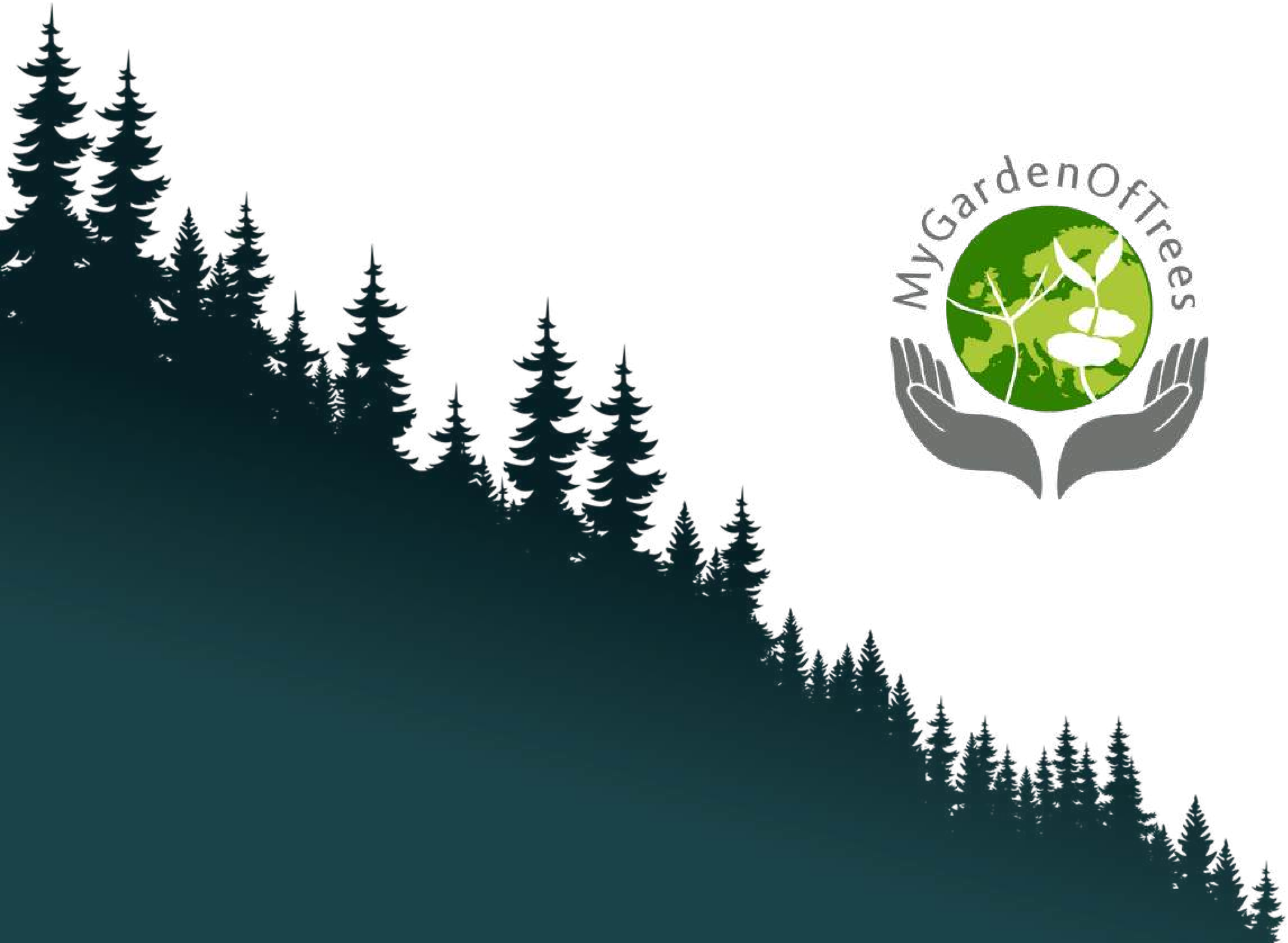

Dear Participant,

You are about to take part in the 2021-2026 trials of MyGardenOfTrees!

We have the pleasure to inform you that during the 2021-2026 trials, 25 participants across Europe will establish their gardens of of different provenances of silver fir (*Abies alba* Mill.) and European beech (*Fagus sylvatica* L.) as well as that of Nordmann fir (*Abies nordmanniana* Spach) from the Caucasus and Oriental beech (*Fagus sylvatica* subsp. *Orientalis* (Lipsky) Greuter & Burdet) from Iran. You are one of these participants, and this protocol explains how to set up your garden.

The team of MyGardenOfTrees is grateful for your participation!

**Katalin Csilléry**
Project leader
Swiss Federal Institute WSL

**Nicole Ponta**
Project coordinator
Swiss Federal Institute WSL

**Haonan Yang**
IT support
Swiss Federal Institute WSL

**Contact:**

### 1. At home: The first steps

#### Received your parcels?

**Please keep the box with the seeds and tags in a cool and dry place** until you can find the time to set up the experiment. Only the “lighter” box has seeds, don’t hesitate to open a check! This can be your garage, your balcony, or any place where temperature does not go above 10˚C.

#### Install and get to know ODK/GIC Collect

The mobile application ODK Collect will be our main communication tool. It can be downloaded free of charge. We will share with you the specific forms of MyGardenOfTrees using your GoogleDrive associated with your Gmail account. Then, you can load these forms in your ODK Collect. Please bear with us, it’s not that complicated at all!

| | | | |
|---|---|---|---|
| 1.<br>Install ODK Collect from the Google Play store, or<br>GIC Collect in the App Store if you have an iPhone. | | ODK Collect | GIC Collect |
| 2.<br>You will receive an email on your **Gmail account** that info@mygardenoftrees has shared setUpMyGarden.xml with you!<br><br>You don’t need to Open it. | | 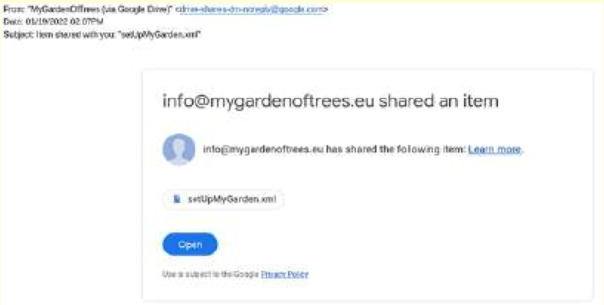 | |
| 3.<br>Open ODK/GIC Collect for the first time at home when you have stable internet connection.<br><br>Choose “Manually enter project details”: | 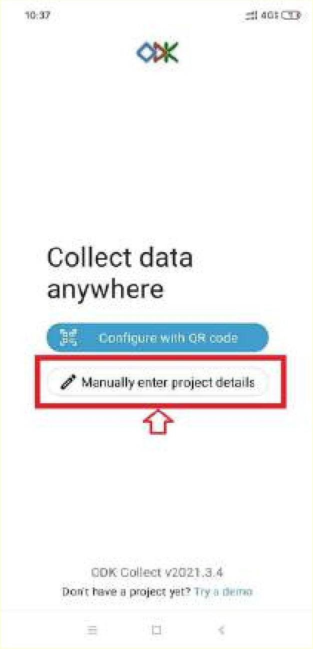 | 4.<br>Choose “Configure” using Google Drive: | 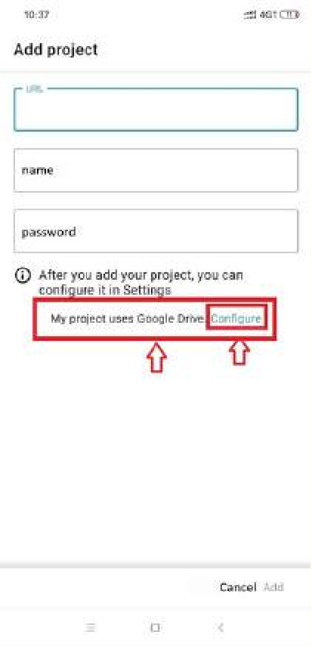 |

5.

Choose “Get Blank Form”:

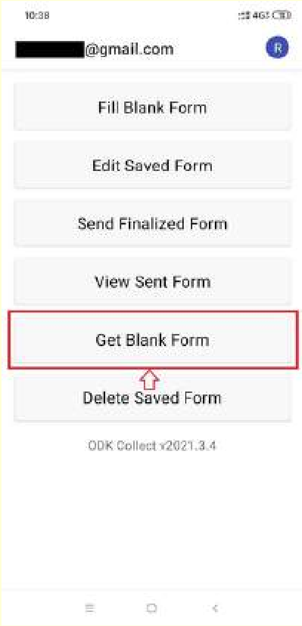


6.

You will see the content of your own Google Drive!

Click “Shared with Me”, and the setUpMyGarden.xml will appear in the list!

Tick the box on the right, and click “Download Selected”!

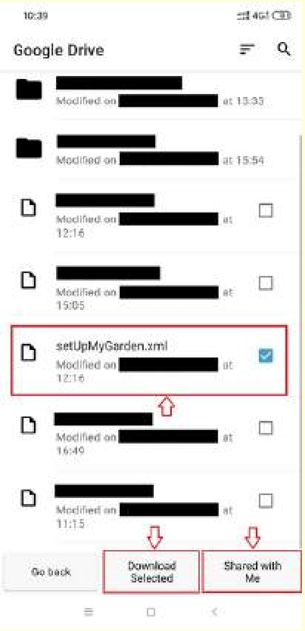


7.

Return to the main menu and click ‘Fill Blank Form’.

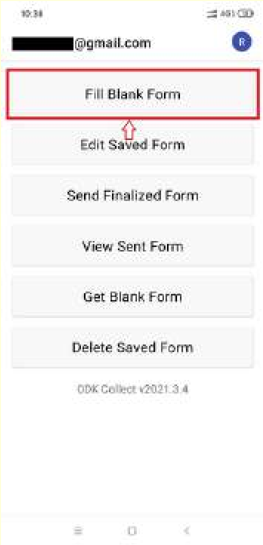


8.

Choose the form SetUpMyGarden!

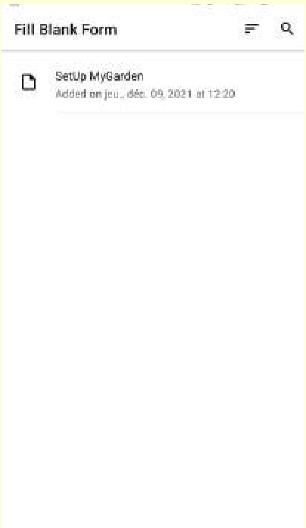


9. The first 3 questions can be filled still in the comfort of your home!

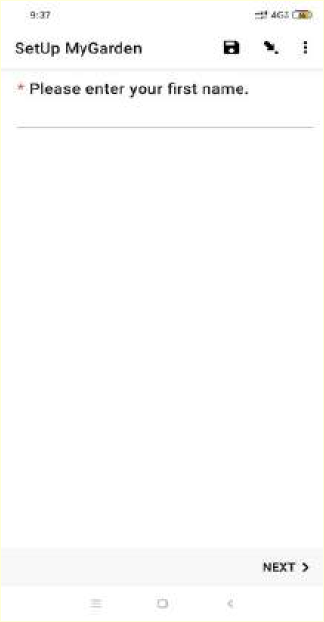


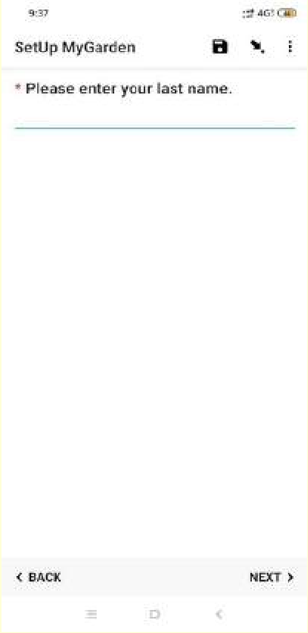


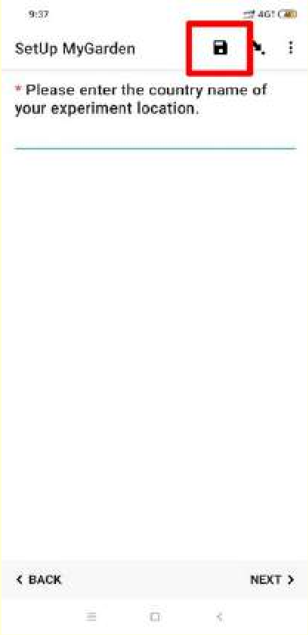


Make sure you **save your form**, so you can continue with your saved form in the forest!

### 2. At home: Preparing to go to the forest

#### The principles

One micro-garden is composed of 4 blocks:

- 2 blocks with *Abies* spp. (called A1 and A2)
- 2 blocks with *Fagus* spp. (called F1 and F2)

Each block has 25 seeding spots. You will sow 10 seeds at each seeding spot.

#### Check the content of your two parcels

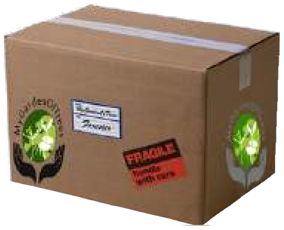

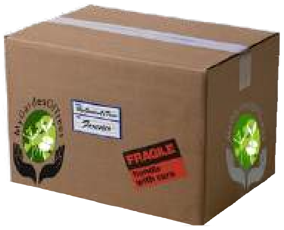

1. 100 plastic tags labeled with species name, provenance and planting location with a bag of seeds attached to them and labeled with species name and provenance
2. A printed list of the species and provenance per block
3. 100 metal seeds protectors
4. 300 metal pegs
5. 4 laminated warning signs (one for each block)
6. A yellow string with knots every meter
7. A squared white sheet of 1m by 1m

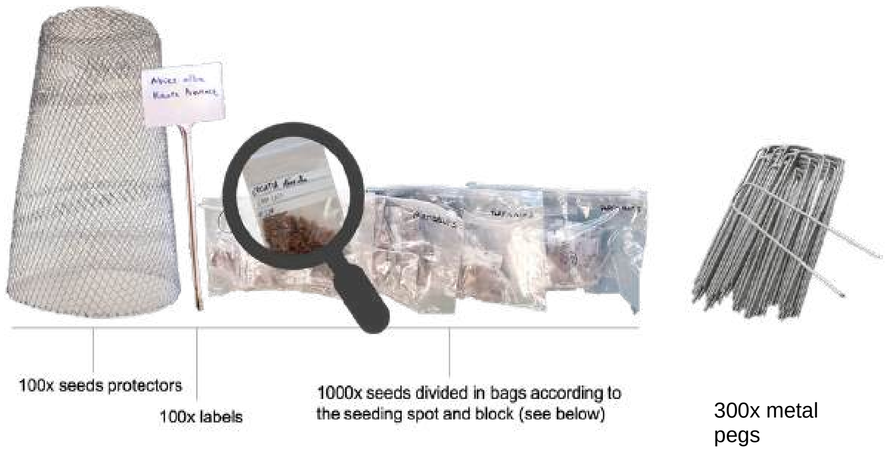


#### What else do you need for setting up the garden?

1. A pair of gardening gloves (to manipulate the seeds protectors!)
2. A hand fork or small rake (to loosen the potentially frozen soil.)
3. A hammer (to hammer down the pegs)
4. A permanent marker to add your contact details to the warning signs, and at least 4 nails to fix them on a tree.
5. Warm clothes and a plastic sheet/pillow to keep warm your knees while sowing!

### Watch this video to get an idea about the work!

(The link is also available on www.mygardenoftrees.eu!)

### 3. In the forest: Setting up your garden

**We recommend that you choose a day with nice weather and, if possible, go with someone else – it's faster and more fun than working by yourself!**

#### Setting up your first block: block F1

We recommend that you start with block F1 (F for Fagus) because Fagus seeds are bigger and slightly easier to sow.

**Materials:** Bring the following material to the selected, roughly 25 m$^2$ large area:

- 25 seeds protectors
- 75 metal pegs (3 for each seed protector)
- plastic tags with seeds for block F1
- the white sheet
- the yellow string
- one laminated sign
- your own tools

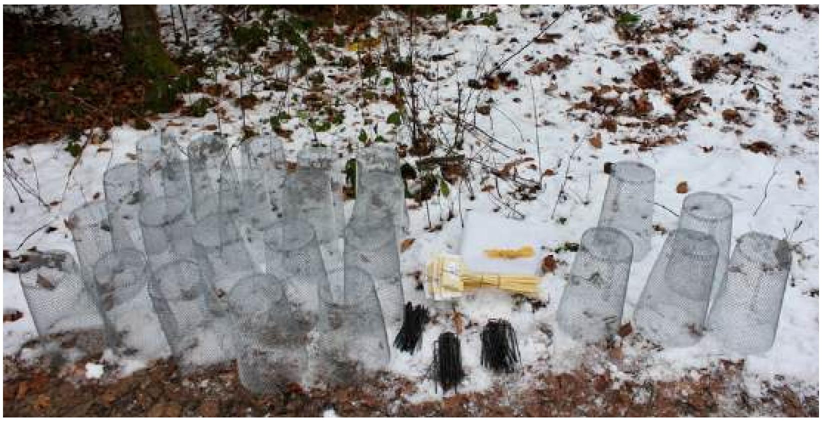

**Clean-up:** Remove fallen branches and other large obstacles from the area.

1.

Open ODK/GIC Collect and choose "Edit Saved Form"

You can continue filling in the same form

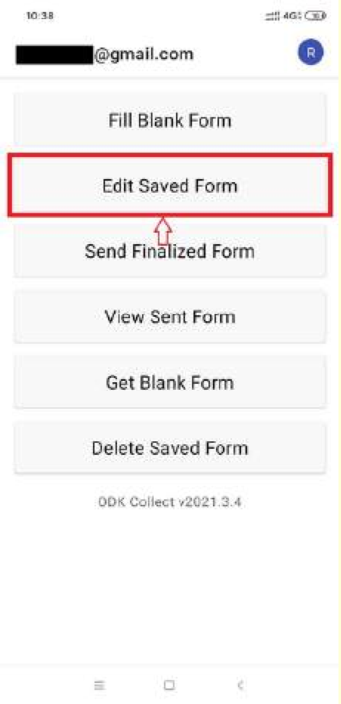


2.

Choose "Go To Start", so you can continue with your form.

You can also modify any questions you have pre-viously answered!

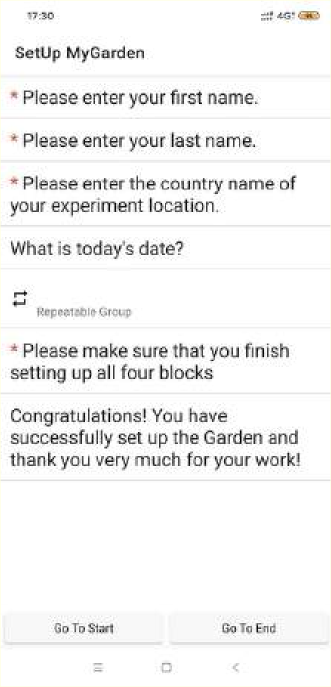

3. Choose block F1!

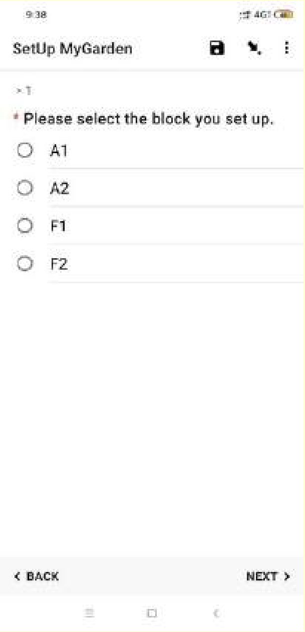


4. Identify the MAGIC corner!

The North-West corner of the area is where the numbering of the seedlings will being. You can use the compass of your phone to identify this corner. Click “Record bearing” and turn your phone till you see NW.

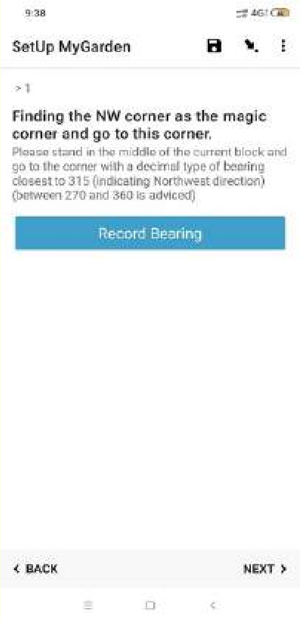


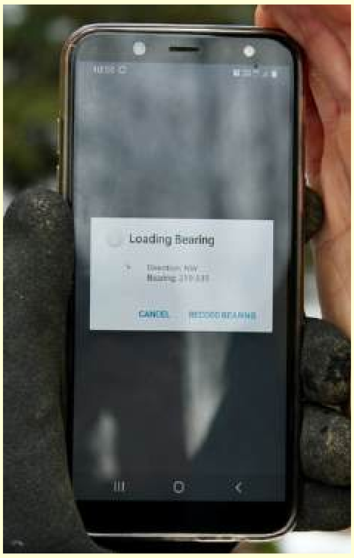


5. Record the GPS coordinates of your block at the MAGIC corner!

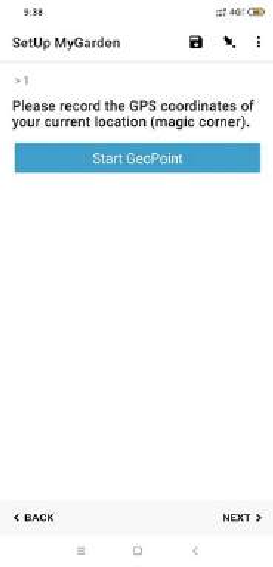


6. Fix a metal peg to the MAGIC corner with the yellow string attached to it and stretch it along the North.

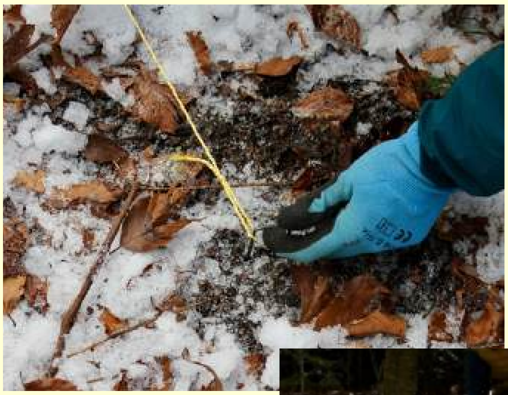

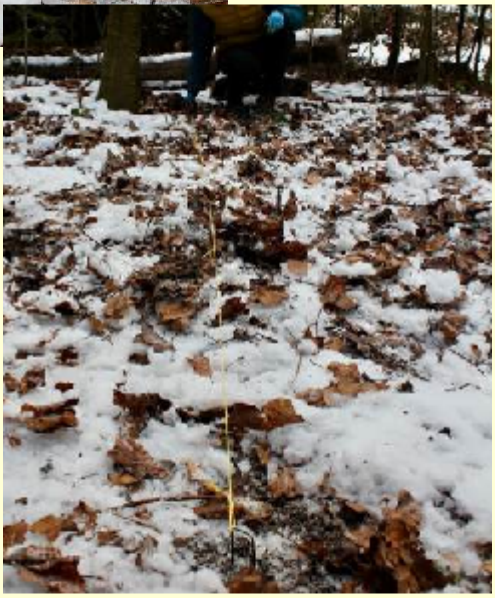

7.

Place the first seed protector at the MAGIC corner. Then, place the 1m by 1m white sheet along the yellow string and place other seed protectors at a distance of 1m from one another.

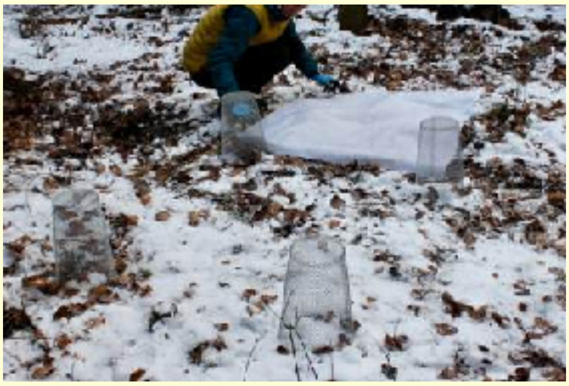

If you encounter an a spot that isn't suitable for sowing or fixing the seed protector to the soil (e.g. roots, stones, etc), skip it and move to the next one!

Continue with a next row until you placed all 25 seed protectors.

You may not be able to establish a perfect 5 by 5 block: it's not a problem, your block can be adapted to the available space.

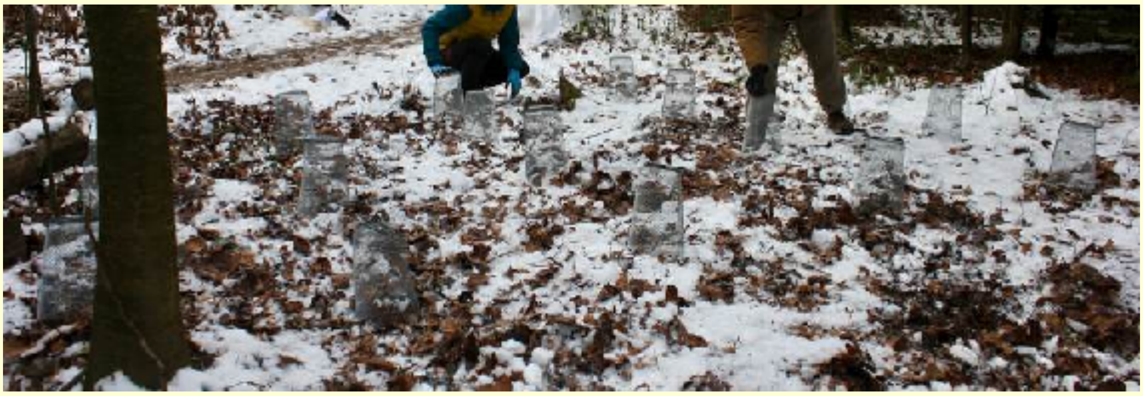

8.

Distribute the labels and seeds, and 3 metal pegs on top of each seed protector starting the numbering in the MAGIC corner and moving towards the North.

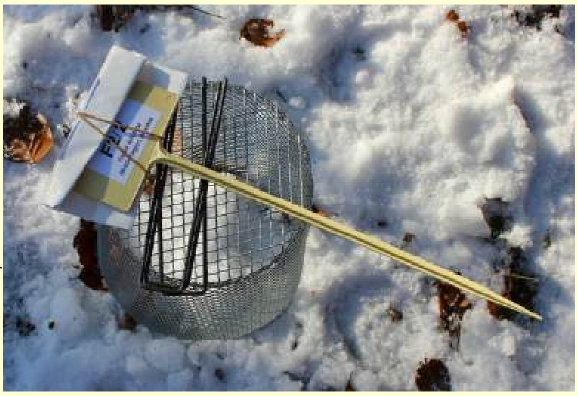

This will help you identify a seedling in later years, in case a label was lost of damaged.

**9.**

**Before you start sowing: Check if the provenance name on the label and on the bag are the same! Although we packed your parcel with great care, errors may still be present! If you detect a mismatch, remove the label from the tag and write by hand the provenance name that is on the seed bag and inform us!**

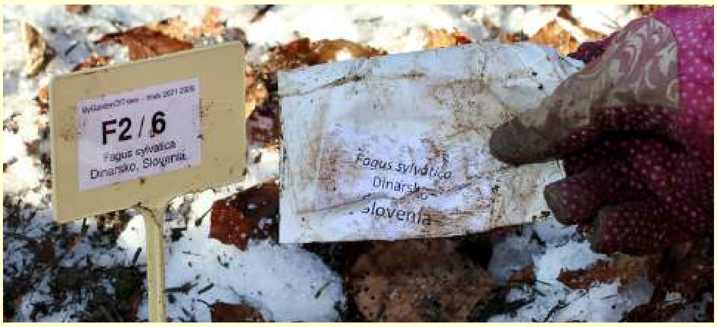

10. Sowing.

- Clean the seeding spot from litter and loosen the soil a bit with a hand fork or a small rake.

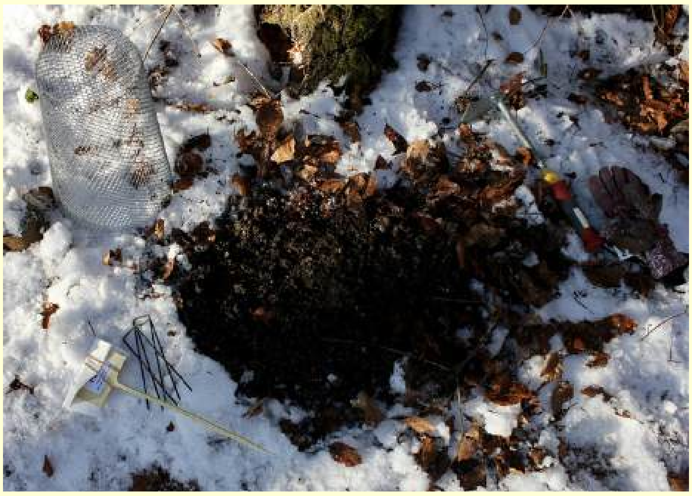

- Place the seeds on the soil, and cover them with a little soil and litter.

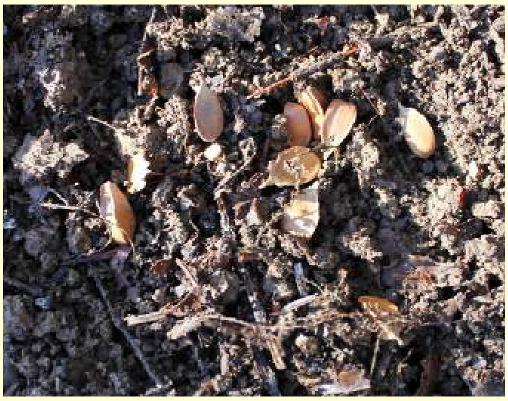

- Place the tag.

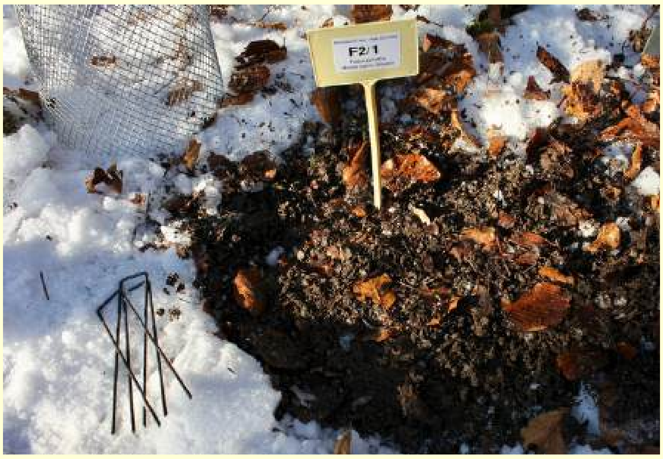


- Place the seed protector, slightly rotate it a coupe of times, and fix it tightly to the soil. Use a hammer to push the pegs well down, but without damaging the seed protector!

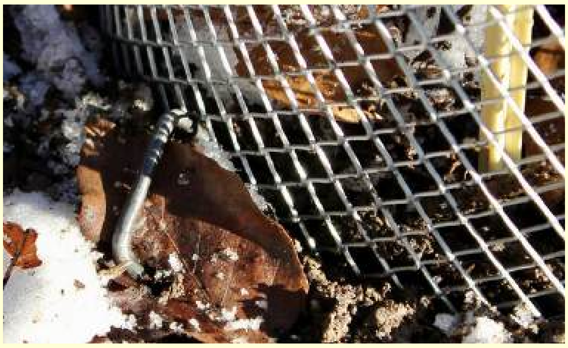

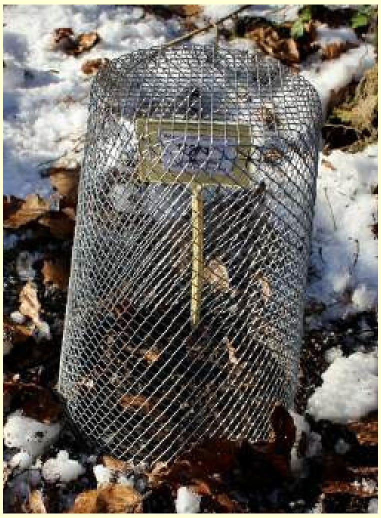

11. Carry on with the other 24 seeding spots and, when you are done, take a picture of your block from the MAGIC corner.

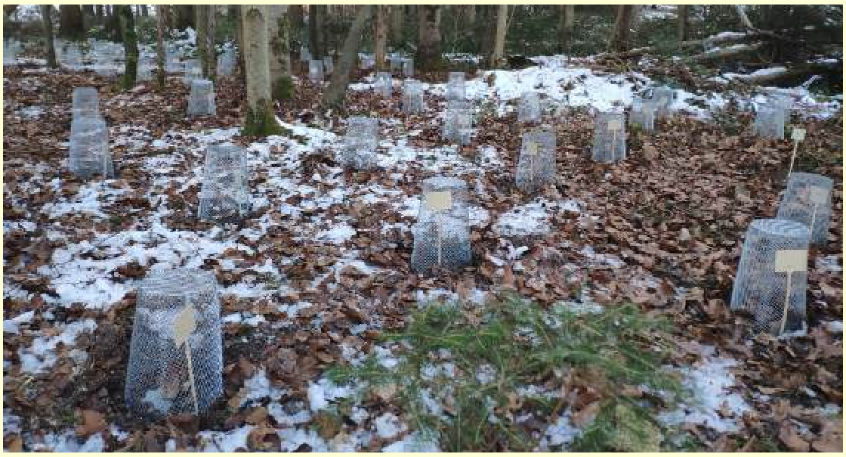

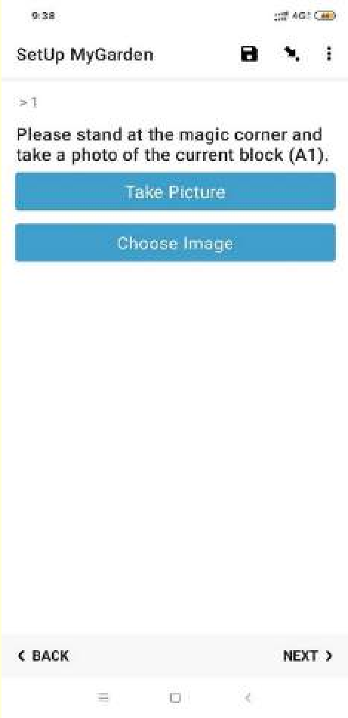


12. Fix the warning sign on a tree!

13. Repeat the same procedure for the 3 other blocks!

It will go faster and faster!

#### Congratulations! Now you have established your garden!

**Make sure you save the ODK Collect form**

**and send it when you have internet connection!**
