## Supplementary figures and images for "Gene–environment interactions govern early regeneration in fir and beech: evidence from participatory provenance trials across Europe"

### A_stage1.jpg

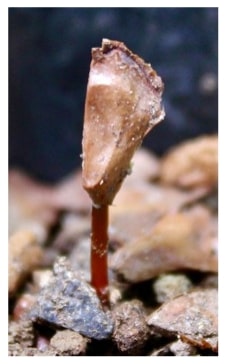

### A_stage1.png

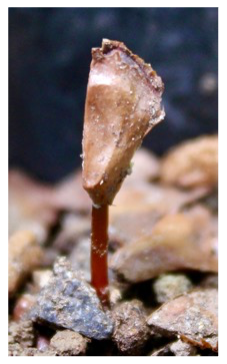

### A_stage2.jpg

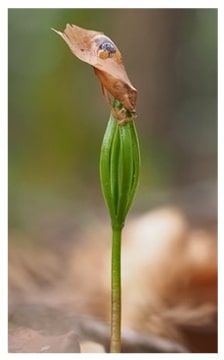

### A_stage2.png

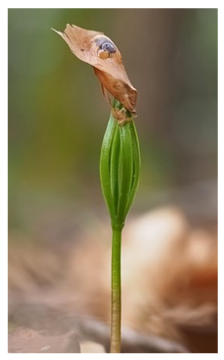

### A_stage3.jpg

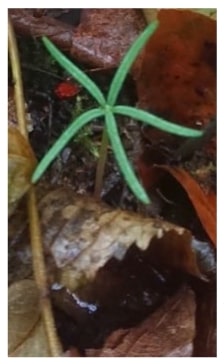

### A_stage3.png

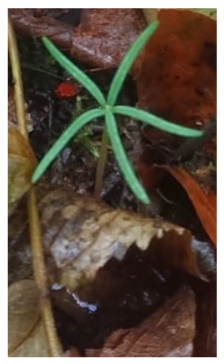

### A_stage4.jpg

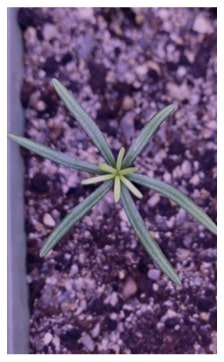

### A_stage4.png

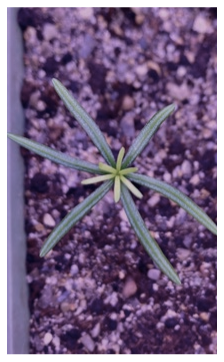

### A_stage5.jpg

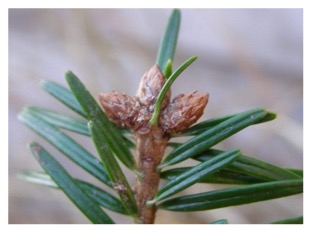

### A_stage5.png

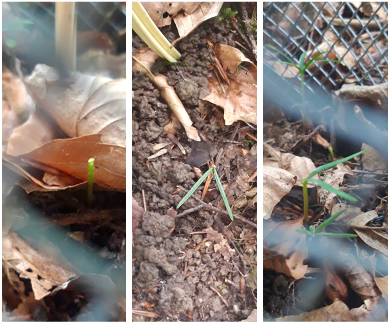

### A_stage6.jpg

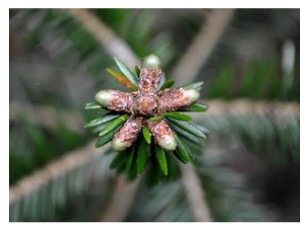

### A_stage7.jpg

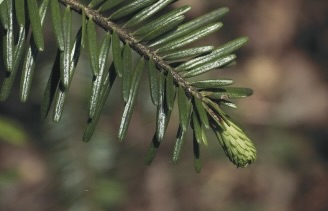

### A_stage8.jpg

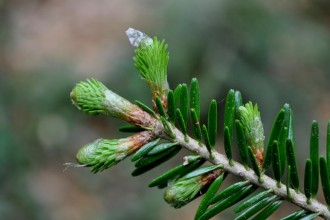

### A_stage9.jpg

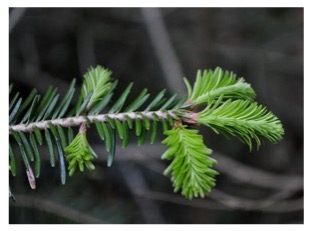

### canopy.jpg

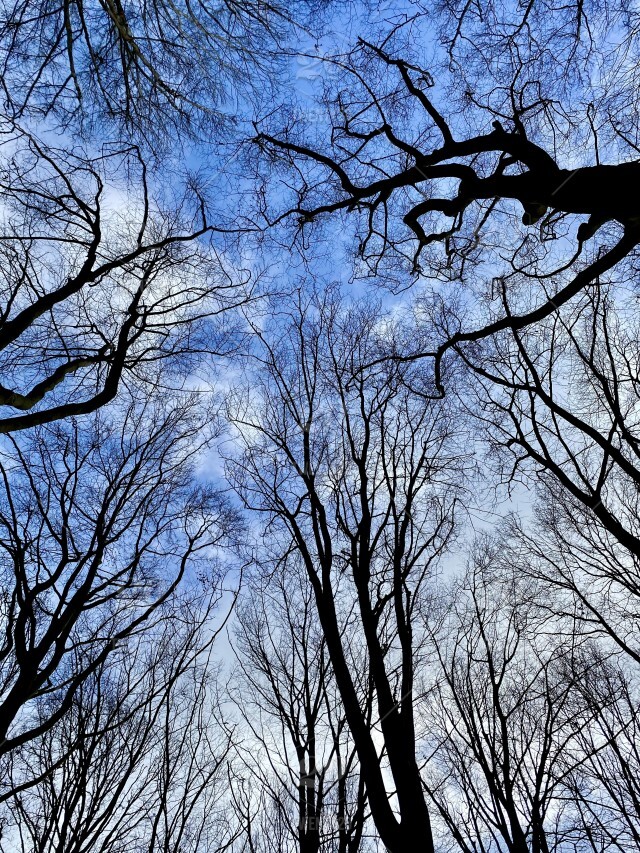

### F_stage1.png

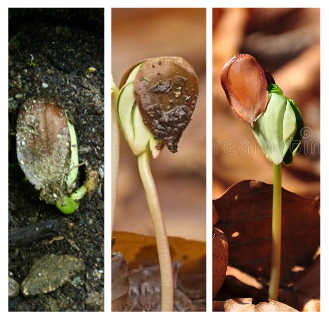

### F_stage2.png

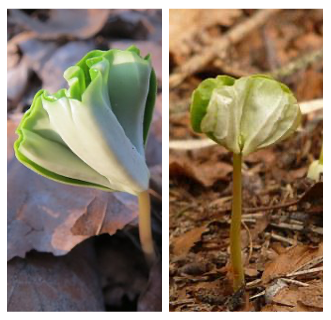

### F_stage3.png

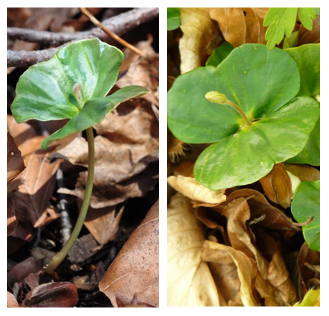

### F_stage4.png

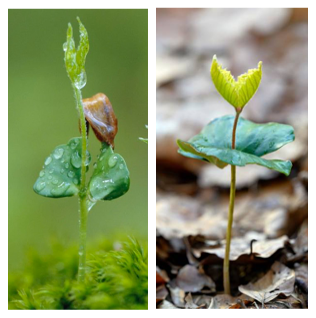

### F_stage5.png

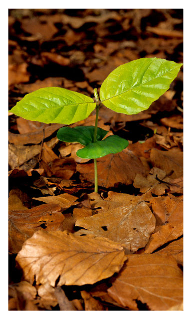

### F_stage6.jpg

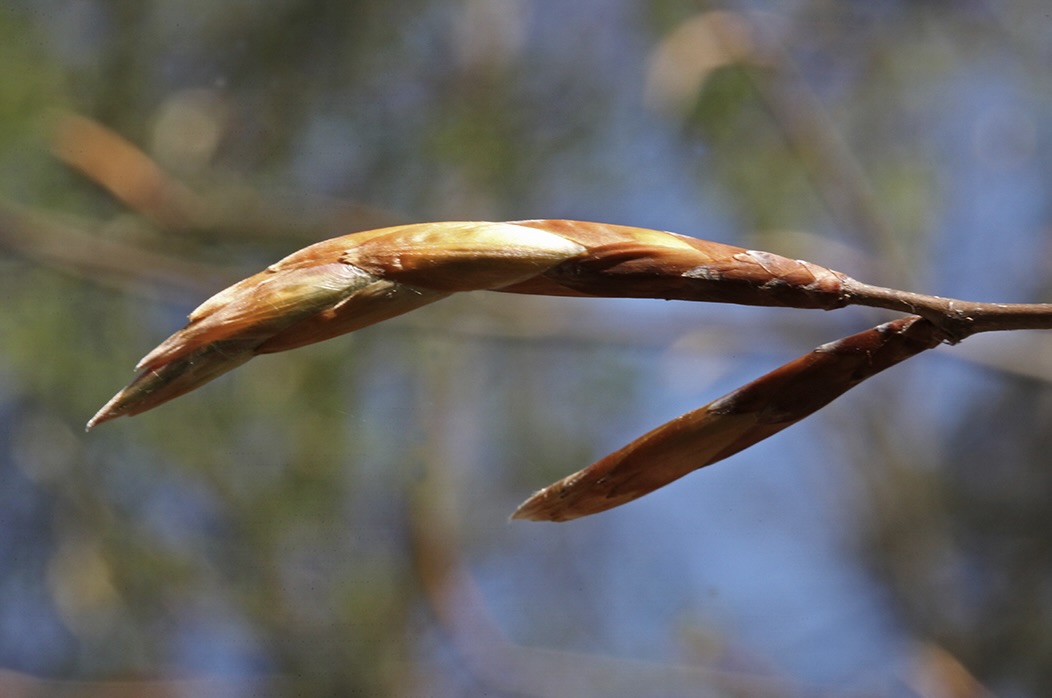

### F_stage6.png

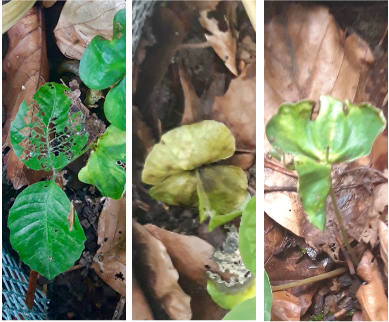

### F_stage7.jpg

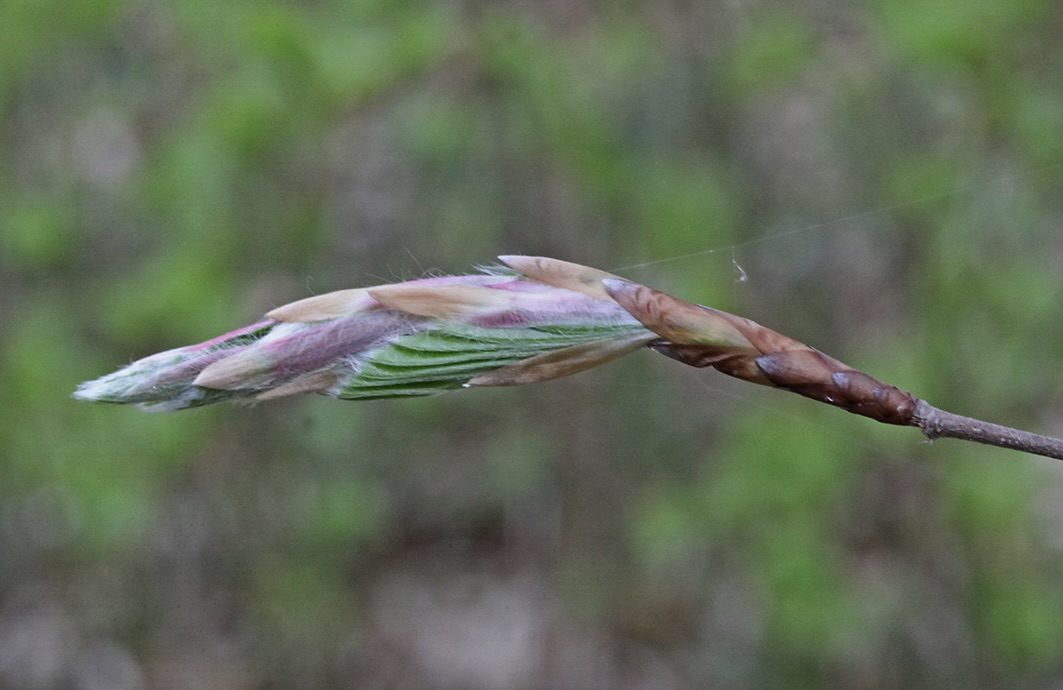

### F_stage8.jpg

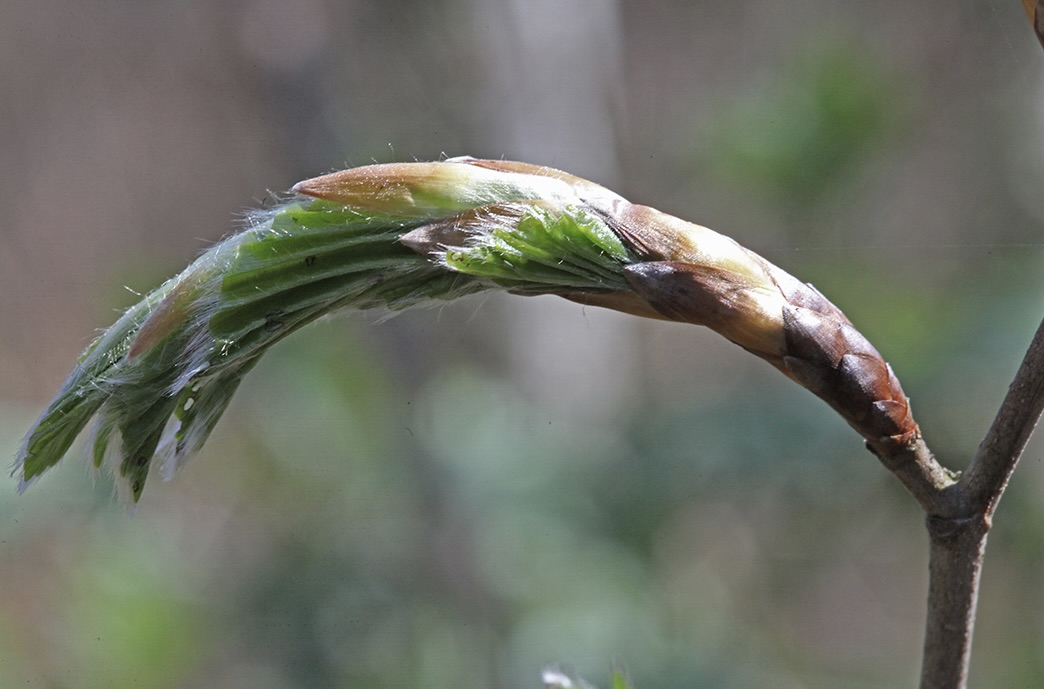

### F_stage9.jpg

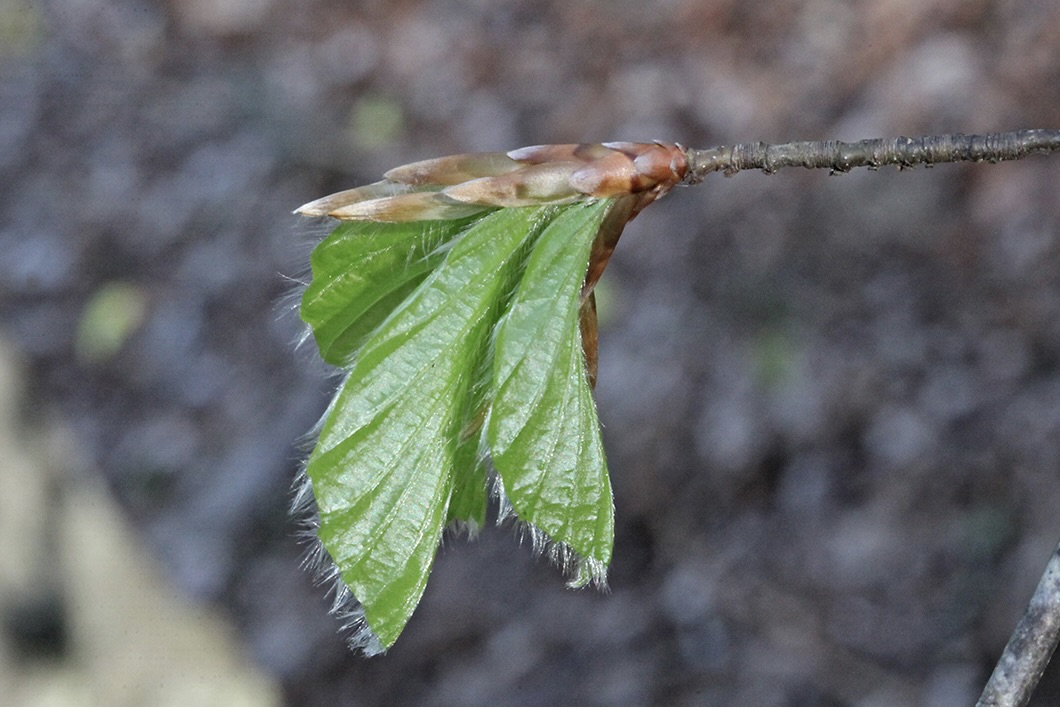

### F_stage10.jpg

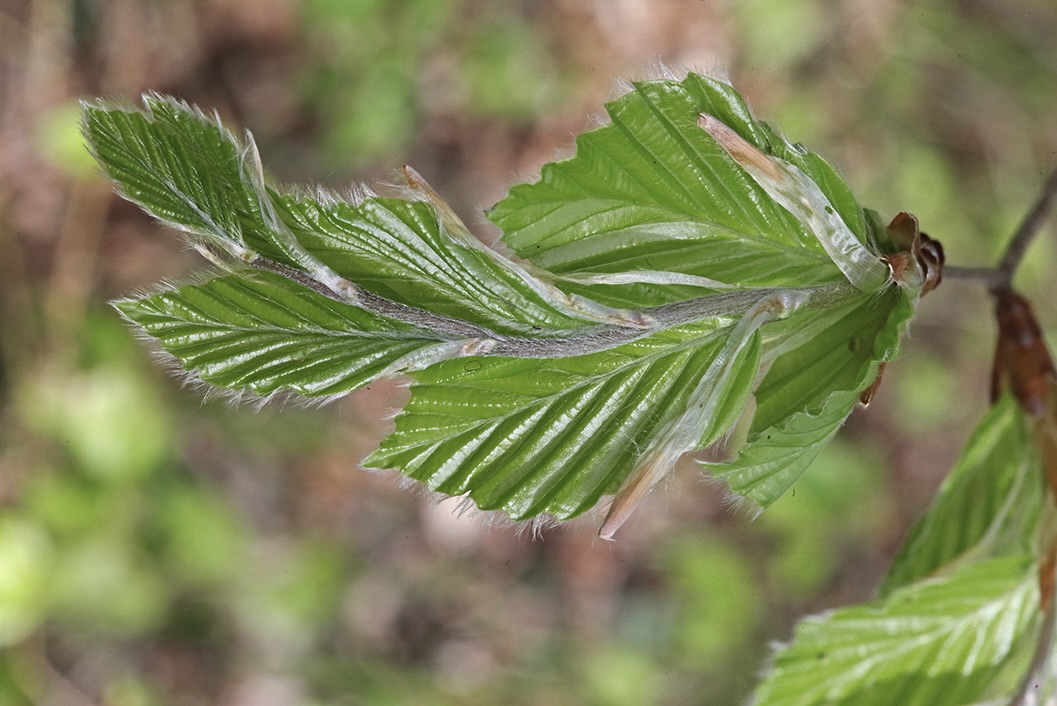

### F_stage11.jpg

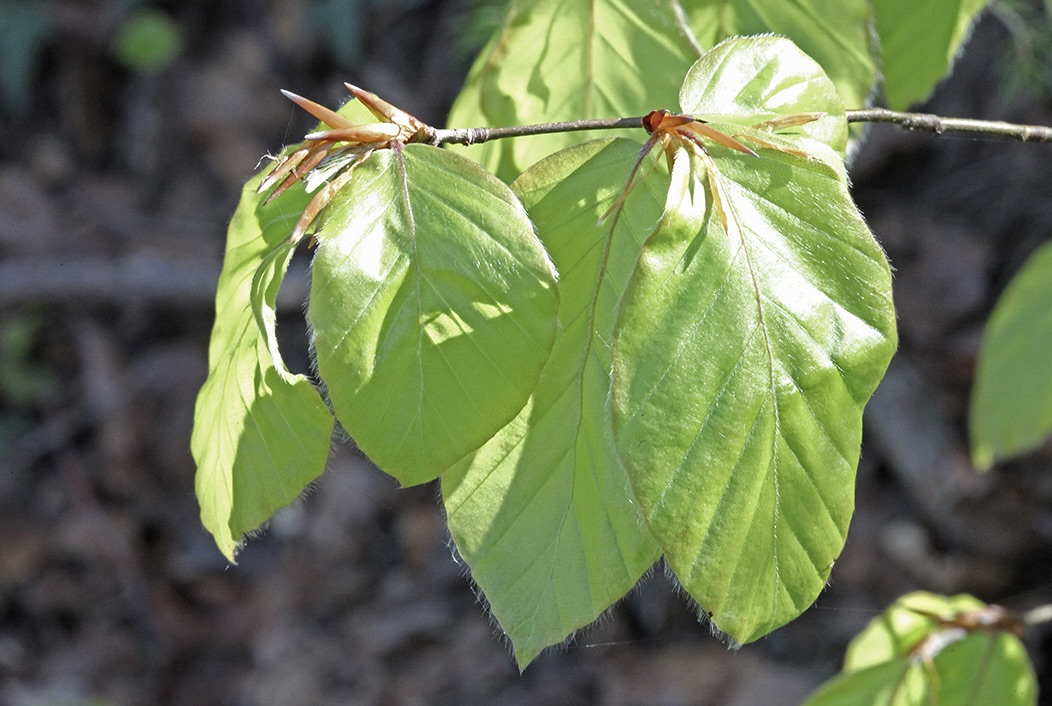

### mesh.jpg

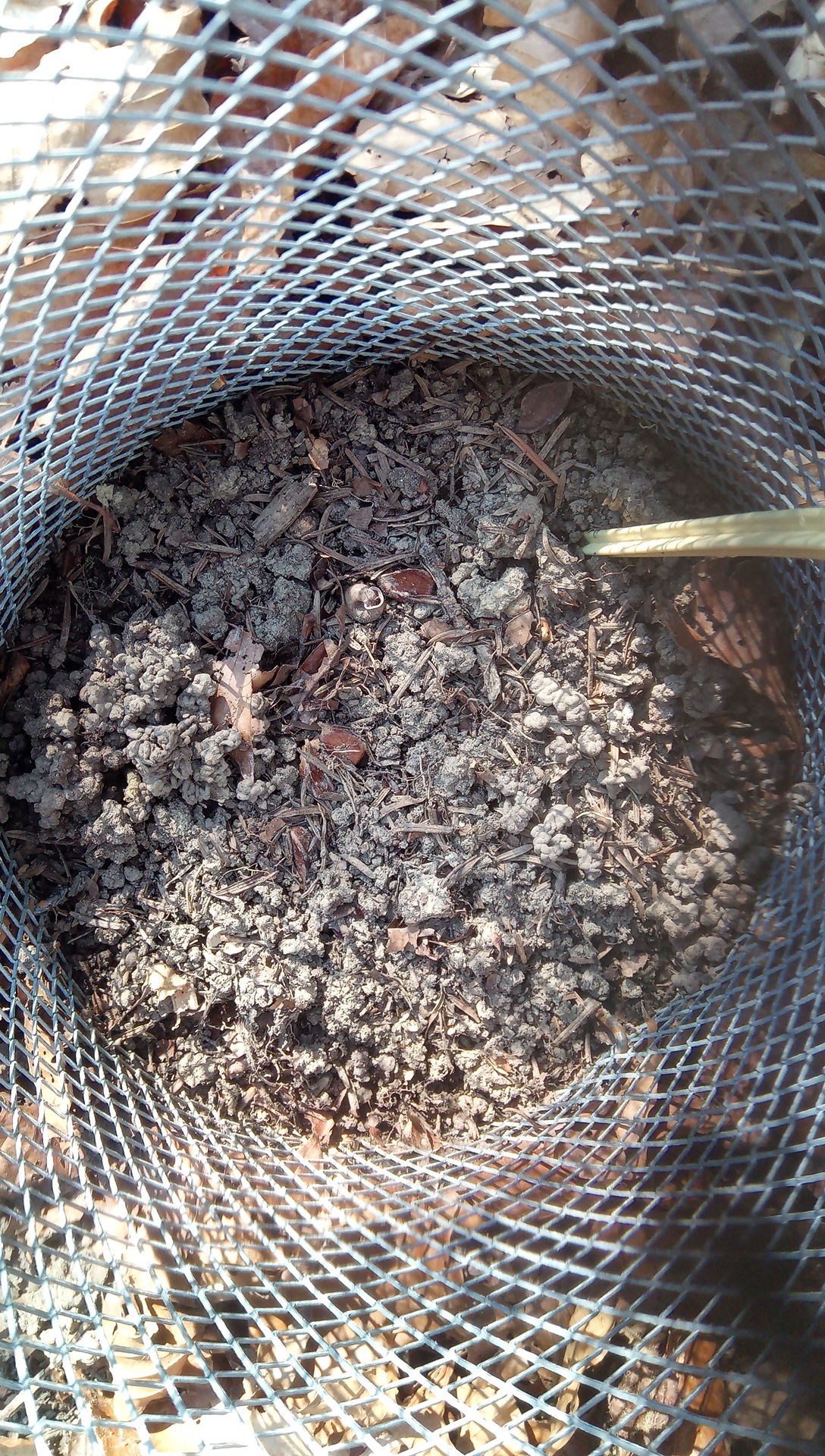
